# Distinct chaperones coordinate the type III secretion system assembly and effector secretion in *Vibrio parahaemolyticus*

**DOI:** 10.64898/2026.11.24.768967

**Authors:** Jananee Jaishankar, Michelle Tan, Kelly A. Servage, Lisa N. Kinch, Kim Orth

## Abstract

Many Gram-negative bacterial pathogens rely on the type III secretion system (T3SS) to deliver virulence factors or effectors into host cells. In *Vibrio parahaemolyticus*, the leading cause of seafood-borne gastroenteritis, one of the two T3SSs, the T3SS2, is essential for intestinal colonization and infection in the human body. While many T3SS2 effectors have been extensively characterized, the chaperones that guide their biogenesis and coordinate assembly of the T3SS machinery remain poorly understood. Here, combining comparative transcriptomic and proteomic approaches with functional analyses, we reveal that four T3SS2 chaperones exert non-overlapping control over distinct clients for pathogenesis. We further show that these chaperones form a hierarchical, multi-layered checkpoint system that ensures the temporal order of secretion, coordinating injectisome assembly, pore formation, and effector deployment. Our findings provide a quantitative framework for understanding the regulation of T3SS2, demonstrating that discrete non-redundant chaperone-mediated assembly is conserved across diverse Gram-negative pathogens.

**Author Summary:** Many bacteria that cause disease use a needle-like molecular syringe, called the type III secretion system, to inject harmful proteins into human cells. Building and operating this syringe requires helper proteins called chaperones, which keep other proteins stable and ready for export. *Vibrio parahaemolyticus*, a leading cause of food poisoning from raw or undercooked seafood, uses one such syringe, T3SS2, to infect the human intestine, yet its chaperones were largely unknown. We identified four chaperones encoded alongside the T3SS2 genes and compared the proteins secreted by normal bacteria and by mutants lacking each one. One chaperone is needed to build the needle itself, a second is needed for the proteins that form a pore in the host cell membrane, and two others each handle different injected toxins, called effectors. Bacteria missing the needle or pore chaperones could not invade intestinal cells, and bacteria missing one of the effector chaperones grew poorly inside host cells. Together, our results suggest that these chaperones act at successive stages to help order the building and use of the syringe. Because this machinery is required for disease, these chaperones may be useful targets for future therapies.

## Introduction

The Type III Secretion System (T3SS) is a complex nanomachine that translocates effectors from the bacterial cytoplasm directly into the cytosol of host cells. The T3SS is a crucial system encoded by many Gram-negative pathogens to mediate disease, including *Yersinia* species, *Salmonella enterica* serovar *Typhimurium*, pathogenic *Escherichia coli*, *Shigella* species, *Bordetella* species, *Pseudomonas aeruginosa*, *Burkholderia pseudomallei*, and *Vibrio* species (1). Also known as an injectisome, the T3SS is a multi-protein complex composed of approximately 20 individual proteins (2). The protein components of the system assemble into a needle-like structure that spans from the cytoplasm through the inner and outer membrane of Gram-negative bacteria (3–7). This needle complex is connected to translocator proteins that, during infection, form a channel in the eukaryotic plasma membrane, facilitating the translocation of effectors directly into the host cell (8). The secretion process is divided into three distinct assembly phases: 1) early substrate secretion, which includes the needle and tip proteins; 2) middle substrate secretion, including the translocon proteins that form the pores in host cell membranes; 3) late substrate secretion for the effectors that are ultimately delivered into the host cytoplasm (9).

Each of the three T3SS assembly phases are coupled to a dedicated set of chaperones that keep the substrates competent for secretion and help establish their hierarchy (9–11) . The chaperones are divided into classes (class I-V) based on their structure and client specificity (12). The early substrate secretion chaperones (class III) prevent the premature cytoplasmic assembly of aggregation prone needle components. The class III chaperones adopt a tetratricopeptide-like repeat (TPR) fold that recognizes the needle substrate as a compex with a small helical co-chaperone (class V) (13, 14). The middle substrate chaperones (class II) also adopt a TPR fold and assist in the secretion of the translocator components that traverse the host cell (15). Finally, class I chaperones bind the late substrate effectors bound for secretion into host cells and include class IA chaperones that bind to a single substrate, and IB chaperones that bind multiple substrates. Despite sharing low sequence similarity, these effector chaperones adopt a conserved structure architecture comprised of an αβα sandwich fold that binds substrate as a dimer (16).

T3SS chaperones are increasingly recognized as important determinants of substrate targeting and the temporal hierarchy of T3SS secretion, in addition to their established roles in substrate stabilization (9). Despite extensive characterization of T3SS regulators and chaperones in many Gram-negative bacteria, identification and functional characterization of these factors in *Vibrio parahaemolyticus (V. para)* remain understudied. *V. para* is a marine pathogen, which causes acute gastroenteritis in humans due to consumption of contaminated seafood (17). The pathogenicity of this bacterium is largely attributed to two T3SSs, one encoded on each chromosome, T3SS1 and T3SS2. The T3SS2 is a major contributor to the enterotoxicity correlated with human disease and observed in animal models (18) and is encoded on the Vp-PAI7 pathogenicity island that is restricted to the O3:K6 pandemic serotypes and related isolates (19).

The Vp-PAI7 pathogenicity island consists of 84 genes. Many of these encode for structural components of the T3SS2 machinery and have been annotated based on sequence homology to other bacterial pathogens (20). In addition, at least ten effectors secreted by T3SS2 have been reported to date (21–29). However, several protein-encoding genes within the island remain unannotated or poorly characterized, leaving gaps in our understanding of how the T3SS2 is regulated and coordinated. To address such gaps, we employed an RNA-sequencing (RNA-seq) methodology to profile highly expressed genes in *V. para* under T3SS2-inducing conditions compared to non-inducing conditions. Four selected candidate genes that were previously uncharacterized and highly upregulated upon T3SS2 induction were then mutated to determine their effect on secretion. Mass spectrometry analysis of secreted proteins from wild-type and mutant strains revealed roles for the candidate genes in orchestrating the secretion of T3SS2 substrates, and further characterization suggests roles for each as chaperones for distinct T3SS2 substrates. Our results indicate that at least four chaperones coordinate the secretion of T3SS2 early, middle, and late substrates in *V. para*. Using genetic and biochemical methods to investigate the interaction between these chaperones and their respective substrates, we were able to elucidate their specificity and potential roles in T3SS2 assembly, effector secretion, and pathogenesis.

## Results

### RNA-seq-based identification of uncharacterized T3SS2 chaperone candidates

The expression of genes within the Vp-PAI7 cluster of *V. para* is governed by a master regulator VtrB, (30, 31) which is activated upon sensing bile acids, specifically taurodeoxycholate (TDC), through the co-component receptor VtrA/C (32) (**Fig 1A**). To identify differentially expressed genes regulated by VtrB, we performed RNA-seq analysis of the *V. para* CAB2 strain (a derivative of the clinical isolate RIMD2210633 deleted for hemolysins and regulator of T3SS1 expression) grown in the presence or absence of TDC. Differentially expressed genes were filtered using a log2 fold change cutoff of 1 and p-value threshold of <0.01. Comparison of induced versus uninduced *V. para* samples identified 850 differentially expressed genes, including 285 upregulated and 565 downregulated genes. Among the upregulated genes, those with particularly high expression (log2 fold change > 4) were predominantly encoded within the Vp-PAI7 pathogenicity island (**Fig 1B**). These include the genes encoding the master regulator (*vtrB*-*vpa1348*), components of the secretion apparatus (*vpa1338*, *vpa1339*, *vpa1341*, *vpa1342*, *vpa1349*), and known effectors (*vopZ*-*vpa1336*; *vopV*-*vpa1357*; *vop*L-*vpa1370*).

**Fig 1:**
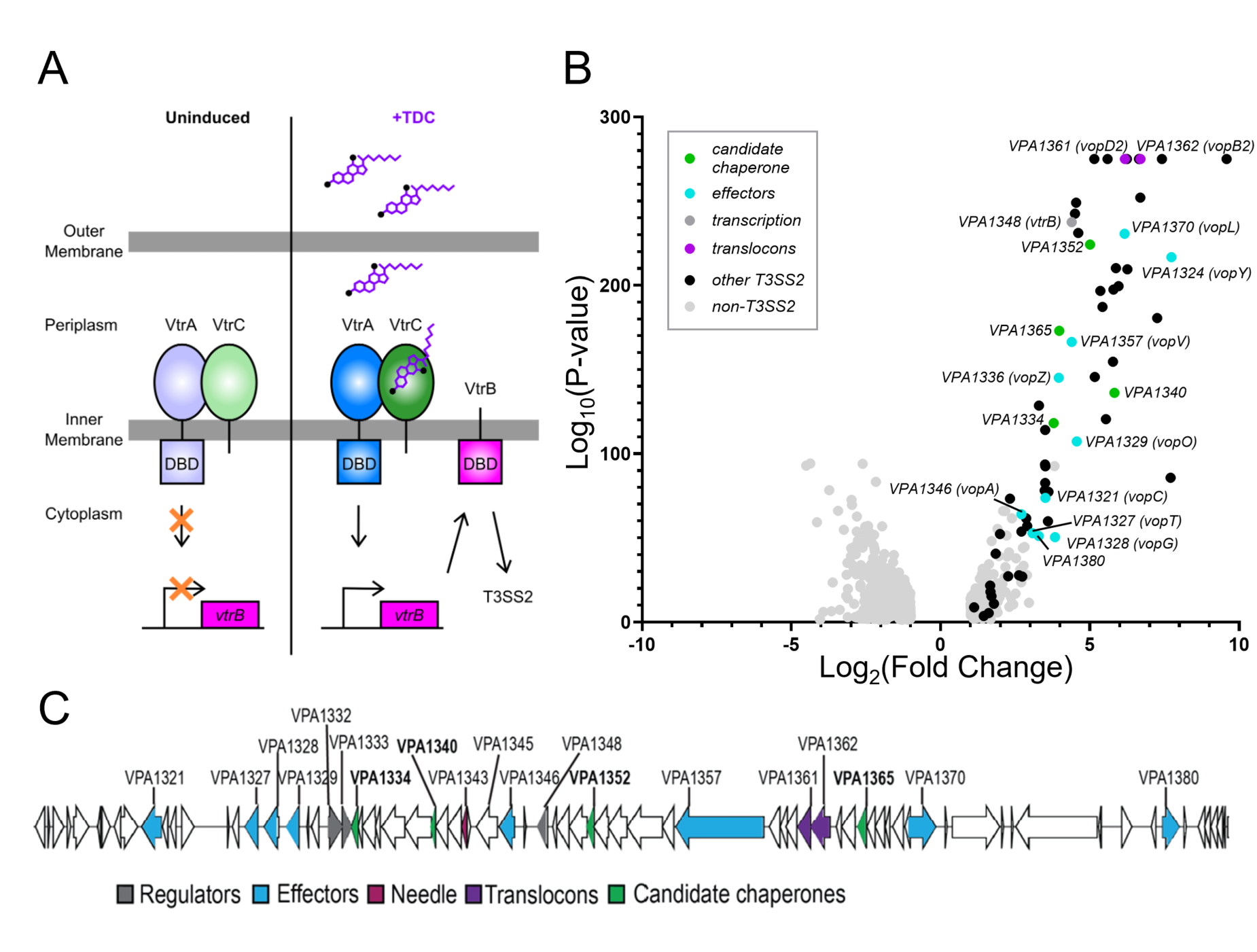
Characterization of differentially expressed *V. para* genes upon bile induction. **(A)** Schematic illustration of T3SS2 induction in *V. para* by the bile acid taurodeoxycholate (TDC). In the absence of bile, the DNA binding domain (DBD) of VtrA does not activate transcription of *vtrB*. Upon TDC binding to the periplasmic domain of VtrC, the DBD of VtrA activates *vtrB* transcription, which subsequently induces the expression of T3SS2 genes (adapted from (30)). (**B)** Volcano plot showing differentially expressed genes in TDC-induced versus uninduced *V. para* CAB2 identified by RNA-seq (p < 0.01, *n* = 3). Genes located within the Vp-PAI7 pathogenicity island are labelled. Genes encoding candidate chaperone-encoding genes are highlighted in green, effectors in cyan, regulators in dark gray, translocons in purple, and other T3SS2 and non-T3SS2 genes in black and light gray, respectively. **(C)** Schematic representation of the Vp-PAI7 pathogenicity island showing T3SS2 genes (arrows), color-coded according to their predicted functions as regulators, effectors, needle protein, translocon proteins, or candidate chaperones, as indicated in the legend.

Several genes were identified with a >10-fold induction upon TDC treatment that have yet to be well characterized. Among these, the candidate genes for this study include *vpa1334*, *vpa1352*, *vpa1365, and vpa1340*. These genes are dispersed across the pathogenicity island and are located at distinct genomic positions relative to the genes encoding T3SS regulators, effectors, translocators, and needle apparatus (**Fig 1C**). Two of these candidate genes encode proteins that are distantly related to class IA/IB effector chaperones by sequence and predicted structure (VPA1334 and VPA1352), while a third (VPA1365) adopts a predicted TPR-like fold found in both class II chaperones responsible for secreting translocators and class III needle chaperones that function with co-chaperones of class V (13–16). While the VPA1340 protein is small and acidic similar to other T3SS chaperones, it does not resemble any of the known classes. Interestingly, as these four candidate genes are situated near multiple genes that are characterized components of T3SS2, we may be able to infer their function by this association.

### Deletion of candidate genes alters *V. para* T3SS2 protein secretion

Using a reverse genetics approach, we generated a panel of isogenic mutant strains in the *V. para* CAB2 strain (22) to define the functional contribution of each candidate gene to T3SS2 expression. We then employed mass spectrometry (MS) to analyze secretomes from these deletion strains. Bacterial cultures of *V. para* CAB2 and each mutant strain (Δ*vpa1334*, Δ*vpa1352,* Δ*vpa1365,* and *vpa1340\**) were grown in the presence of TDC and KCl for 3 hours to induce T3SS2 expression and mimic host cell contact, respectively. Cells were then spun down, and culture supernatants were collected to isolate secreted proteins for analysis by MS.

Comparative analysis of the secretomes from each mutant strain was performed using label-free quantitative Liquid Chromatography-Tandem Mass Spectrometry (LC-MS/MS). Protein abundances from the four mutant strains were each compared to the CAB2 strain in biological triplicate and then visualized using volcano plots (**S1 Fig**). Interestingly, mutation of each candidate gene resulted in both distinct and significant changes in the abundances of many secreted proteins relative to the wild-type strain. For the purposes of this study, we focused primarily on secreted proteins exhibiting a significant decrease in abundance in the mutant strains (Fold change < 0.5 and p-value < 0.05), assuming these changes are indicative of disruptions in secretion pathways and would provide insight into the functional roles of the deleted genes. Since each of the four candidate gene deletions produced a distinct proteomic fingerprint, the mutations do not appear to cause a general defect in secretion but instead differentially affect subsets of secreted proteins and support the idea that each of these genes (*vpa1334*, *vpa1352*, *vpa1365, and vpa1340)* play a functional role in the T3SS2-mediated virulence of the bacterium.

### VPA1334 preferentially mediates effector secretion

Examining the quantitative proteomic results of secreted T3SS2 proteins revealed that deletion of *vpa1334* significantly impairs the secretion of two effectors, VPA1321 (VopC) and VPA1370 (VopL) (Fold change < 0.1 relative to *V. para* CAB2 strain) (**Fig 2A**). AlphaFold predictions of the full length effector sequences for VopC and VopL with the VPA1334 dimer indicate relatively poor interactions with ipTM scores of 0.41 and 0.58, respectively. This low confidence value, which scores the interactions based on the whole chain, reflects the presence of the effector domains that do not interact with the chaperone and therefore decrease confidence scores. Using VPA1334 with the N-terminal translocation signal sequence of VopC and VopL produced high confidence interaction scores (ipTM=0.74 and 0.73, respectively) (**Fig 2B, S1 Table**). Both structure models exhibit well-defined interaction interfaces as evidenced by confidence metrics, and the interactions resemble those of known effector-chaperone complex structures with translocation signal sequences. The predictions suggest that VPA1334 interacts with the N-terminal regions of VopC and VopL.

**Fig 2:**
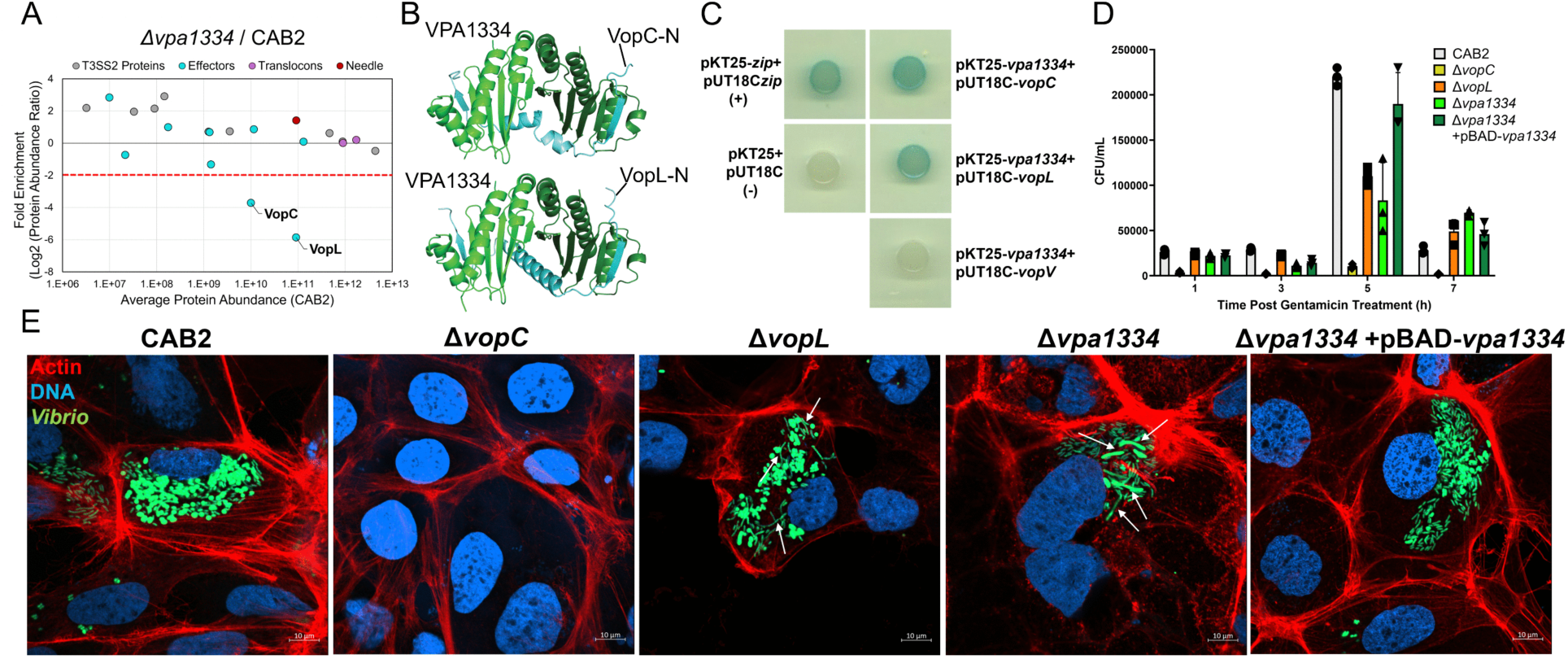
VPA1334 interacts with T3SS2 effectors and contributes to intracellular survival of *V. para*. **(A)** Plot of T3SS2 proteins identified by label-free LC-MS/MS comparing Δ*vpa1334* and CAB2 secreted protein fractions. Fold enrichment (Δ*vpa1334*/CAB2) is plotted against the average protein abundance in the CAB2 strain (*n = 3*). T3SS2 effectors are highlighted in cyan, the needle protein in magenta, and the translocons in purple. The red horizontal dotted line indicates the significance threshold (Log2(Δ*vpa1334*/CAB2) < -2). **(B)** Structure showing the AlphaFold-predicted interaction between VPA1334 (green) and the N-terminal translocation sequence of VopC or VopL (cyan): iPTM scores of 0.74 (VopC) and 0.73 (VopL). **(C)** Bacterial two-hybrid assay demonstrating interactions between VPA1334 and its putative binding partners. Positive interactions between VPA1334 and VopC, and between VPA1334 and VopL, are indicated by blue colonies. No interaction was detected between VPA1334 and the effector VopV, as indicated by white colonies. The negative control (pKT25 + pUT18C) and positive control (pKT25-zip + pUT18C-zip) were included for comparison. **(D)** Intracellular survival of the indicated *V. para* strains in Caco-2 cells. Colony-forming units (CFU) were quantified following 1.5 h of infection, removal of extracellular bacteria by gentamicin treatment for 1.5 h, and enumeration at 1, 3, 5, and 7 h post-gentamicin treatment (PGT). **(E)** Representative confocal micrographs of Caco-2 cells at 5 h PGT infected with GFP-expressing intracellular *V. para* (green). Filamentous bacteria are indicated by white arrows. Actin was stained with rhodamine-phalloidin (red) and DNA was stained with Hoechst (blue).

To probe whether VPA1334 forms a stable complex with VopC or VopL we employed the Bacterial Adenylate Cyclase-Based Two-Hybrid (BACTH) system. Full-length *vpa1334* was fused with the T25 fragment of *cyaA*, while full-length *vopC* or *vopL* were fused to the T18 fragment of *cyaA*. Co-transformation of these constructs into *E. coli* BTH101 resulted in the formation of blue colonies, indicating a physical interaction between VPA1334 and both VopC or VopL (**Fig 2C**). As a negative control, we tested the interaction of VPA1334 with another effector, VopV. In this case, the absence of blue colonies indicated no interaction with VPA1334, suggesting that VPA1334 specifically associates with VopC and VopL and may function as a chaperone for these two effectors.

We next wanted to identify the effect of *vpa1334* deletion on the ability of CAB2 to invade and replicate within the colonic epithelial Caco-2 cell line. We performed a gentamicin protection assay using *V. para* strains CAB2, Δ*vpa1334*, *ΔvopC, ΔvopL*, and the complementation strain for Δ*vpa1334 (Δvpa1334+*pBAD*-vpa1334)* along with accompanying confocal images of invaded Caco-2 cells at 5h post gentamicin treatment (PGT). For the *V. para* CAB2 strain, we observed the expected profile for T3SS2-mediated invasion (**Fig 2D**). As observed previously, deletion of *vopC* inhibits invasion of *V. para* completely while deletion of *vopL* results in an invasion defect as evidenced by a reduction in intracellular bacterial growth and many stressed, filamented bacteria (**Fig 2D,E**) (23). Deletion of *vpa1334* produces a similar intracellular growth profile as the *vopL* deletion mutant with reduced intracellular bacteria and a filamentous morphology inside host cells (**Fig 2D,E**). Complementation of VPA1334 on a plasmid (*Δvpa1334+*pBAD*-vpa1334*) corrected these invasion defects, confirming that the observed phenotype was specifically due to the absence of VPA1334. These observations further support the conclusion that VPA1334 might act as an essential chaperone for VopL and VopC playing a critical role in promoting the intracellular survival of *V. para* in Caco-2 cells.

### VPA1352 defines a non-redundant pathway for effector secretion

Since our data supports VPA1334 functioning as a chaperone, we tested VPA1352, another candidate class 1 chaperone that was upregulated in our RNA-seq screen, using the same secretome profiling approach. Proteomic data of T3SS2 secreted proteins showed that deletion of *vpa1352* resulted in significant reduction of the effector VopV (**Fig 3A**). Similar to the AlphaFold predictions for VPA1334, the full length VopV effector sequence modeled with the VPA1352 dimer indicate a relatively poor interaction(ipTM score of 0.44), reflecting the high content of repeats and disordered sequence encoded by the large VopV protein (1622 amino acid residues long, with a low pTM of 0.28). Modeling VPA1352 with the N-terminal translocation signal sequence of VopV produced a high confidence interaction score (ipTM=0.78). (**Fig 3B, S1 Table**). To test binding, we employed the BACTH assay where full-length VPA1352 was fused to the T25 fragment of cyaA in pKT25, and the coding sequence corresponding to amino acids 1-200 of VopV was fused to the T18 fragment in pUT18C. Co-transformation of these constructs into *E. coli* BTH101 yielded blue colonies, confirming a physical interaction between VPA1352 and VopV(1-200aa) (**Fig 3C**). To validate whether this interaction is specific for the effector VopV, we performed a control BACTH assay with VPA1352 and the effector VopL and observed no blue colonies. While VopV is known to be a critical factor for bacterial enterotoxicity (21), its contribution to bacterial invasion and intracellular replication is unexplored. We performed a gentamicin protection assay of Caco-2 cells infected with Δ*vopV*, Δ*vpa1352*, and Δ*vpa1352+*pBAD*-vpa1352* strains to compare with the CAB2 strain. Results showed that deletion of *vopV* or *vpa1352* did not affect intracellular replication of *V. para* in Caco-2 cells (**Fig 3D**). This result was also confirmed by confocal microscopy of invaded Caco-2 cells at 5h PGT (**Fig 3E**). Collectively, our data indicates that both candidate proteins VPA1334 and VPA1352 act as chaperones for three specific effectors in *V. para*.

**Fig 3:**
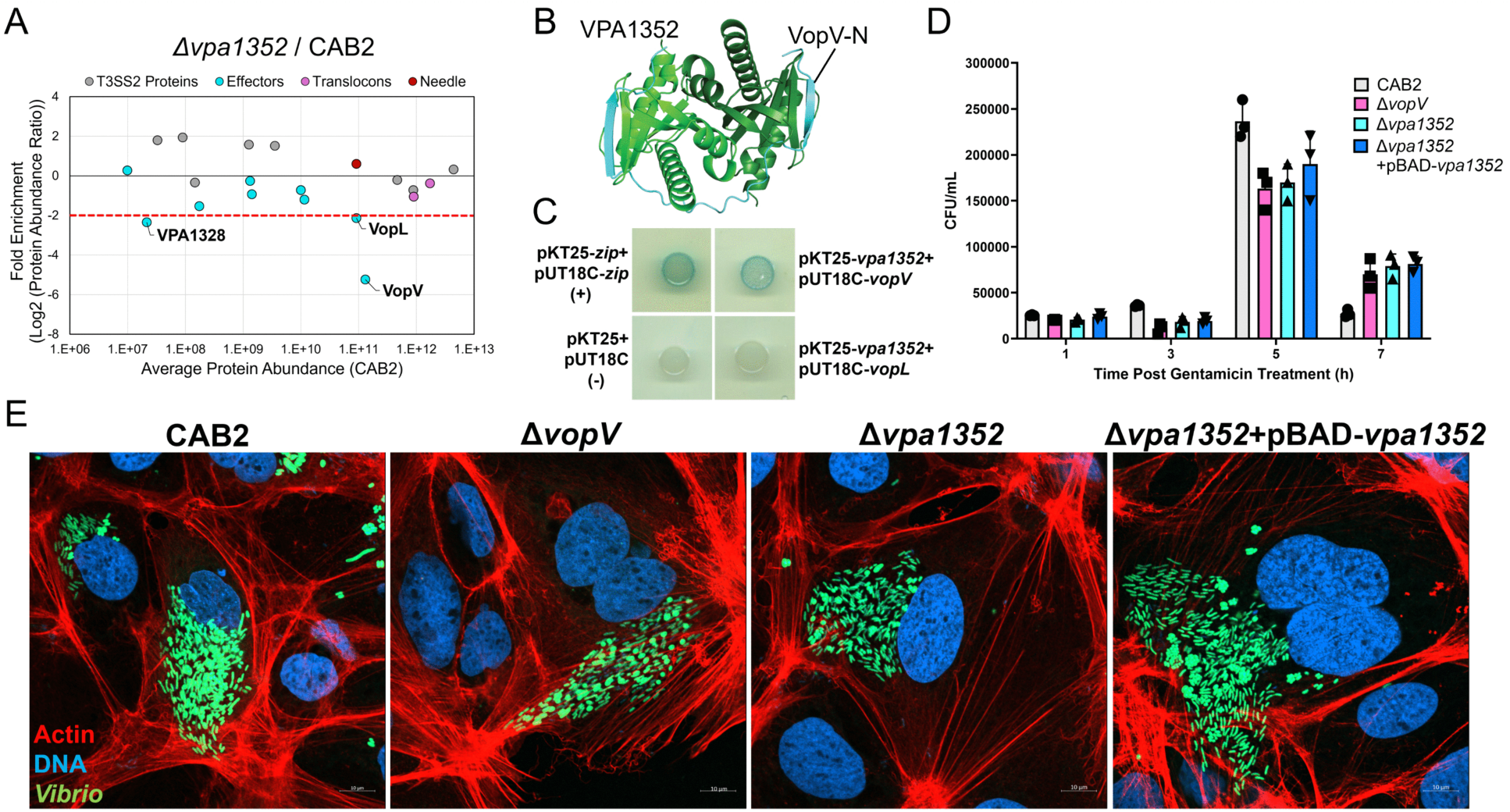
Loss of VPA1352 disrupts a distinct T3SS2 effector secretion pathway. **(A)** Plot of T3SS2 proteins identified by label-free LC-MS/MS comparing Δ*vpa1352* and CAB2 secreted protein fractions. The fold enrichment (Δ*vpa1352*/CAB2) is plotted against the average protein abundance in the CAB2 strain (*n = 3*). T3SS2 effectors are highlighted in cyan, the needle protein in magenta, and the translocons in purple. The red horizontal dotted line indicates the significance threshold (Log2(Δ*vpa1352*/CAB2) < -2). **(B)** Structure showing the AlphaFold-predicted interaction between VPA1352 (green) and the N-terminal region of VopV (amino acids 1–200; cyan) (ipTM=0.78). **(C)** Bacterial two-hybrid assay demonstrating the interaction between VPA1352 and VopV. A positive interaction was observed, as indicated by the formation of blue colonies. No interaction was detected between VPA1352 and the effector VopL, as indicated by white colonies. The negative control (pKT25 + pUT18C) and positive control (pKT25-zip + pUT18C-zip) were included for comparison. **(D)** Intracellular survival of the indicated *V. para* strains in Caco-2 cells. Colony-forming units (CFU) were quantified following 1.5 h of infection, removal of extracellular bacteria by gentamicin treatment for 1.5 h, and enumeration at 1, 3, 5, and 7 h post-gentamicin treatment (PGT). **(E)** Representative confocal micrographs of Caco-2 cells at 5 h PGT infected with GFP-expressing intracellular *V. para* (green). Actin was stained with rhodamine-phalloidin (red) and DNA was stained with Hoechst (blue).

Deletion of each chaperone leads to a decrease in secretion levels of their associated effector substrates; however, it has been proposed that effector chaperones can also affect the stability of their binding partners in the cytosol of bacterial cells (33) . To probe this hypothesis, we used LC-MS/MS to profile protein levels from the cell pellet fractions of *Δvpa1334* and *Δvpa1352* strains to compare to the secreted protein profiles. Results showed no significant changes in the abundance of effectors VopC, VopL, or VopV in the pellet fractions (**S2A/D and B/E Fig**). Since there are at least 7 additional effectors known in *V. para*, we also constructed a double deletion strain lacking genes encoding both chaperones (*Δvpa1334Δvpa1352*) to test whether removal of both chaperones could affect additional effectors. LC-MS/MS analysis of the secreted protein fraction from the double mutant *Δvpa1334Δvpa1352* revealed an additive effect of the individual deletions, showing a significant decrease in abundance of the three expected effectors (VopC, VopL, and VopV) (**S2F Fig**). Notably, only two effectors VPA1328 (VopG) and VPA1329 (VopO) show lowered secretion levels in the double mutant compared to the single deletion mutants. Similar to the single deletion mutants, levels of the effectors were not significantly affected in the cell pellet fraction of the *Δvpa1334Δvpa1352* strain (**S2C Fig**). This result suggests that these chaperones regulate distinct effectors of T3SS2.

### VPA1365 is required for pore formation

In addition to effector-specific chaperones, another uncharacterized protein, VPA1365, was identified as a potential chaperone in our RNA-seq screen for upregulation by TDC. To investigate its potential role in T3SS2 regulation, we generated a deletion mutant of *vpa1365* and employed LC-MS/MS to determine the effect on protein secretion. Interestingly, the secretion of effectors was unimpaired in the *Δvpa1365* strain as compared to the wild-type strain, but there was a severe decrease in levels of VPA1362 (VopB2), a major translocon for *V. para* (**Fig 4A**). To test if VPA1365 acts as a chaperone for VopB2, we first generated AlphaFold models for the interaction. The modeled structure complex for the full-length translocon highlighted an interaction between its N-terminus and the chaperone. Using the N-terminal sequence for the VopB2 translocon to model the complex produced a confident interaction (ipTM=0.8) (**Fig 4B, S1 Table**). However, VPA1361 (VopD2), the minor translocon present in *V. para* is not predicted to bind to VPA1365 based on AlphaFold modeling. To further test translocon binding, we employed a BACTH assay as described previously with *vpa1365* and either *vopD2* or *vopB2*. Co-transformation of these constructs into *E. coli* yielded blue colonies only in the case of pKT25-*vpa1365* and pUT18C-*vopB2*, confirming a physical interaction between VPA1365 and VopB2 (**Fig 4C**). A decrease in translocon secretion would not be expected to affect the secretion of effectors in the absence of host cells, as described previously (34). To confirm this expectation, we used Western blotting to measure the levels of two representative effectors VopL and VopA in the *Δvpa1365* strain. As shown in **Fig S3**, both effectors were detected in the supernatant of the *Δvpa1365* strain equivalent to the amount secreted for the wild-type strain.

**Fig 4:**
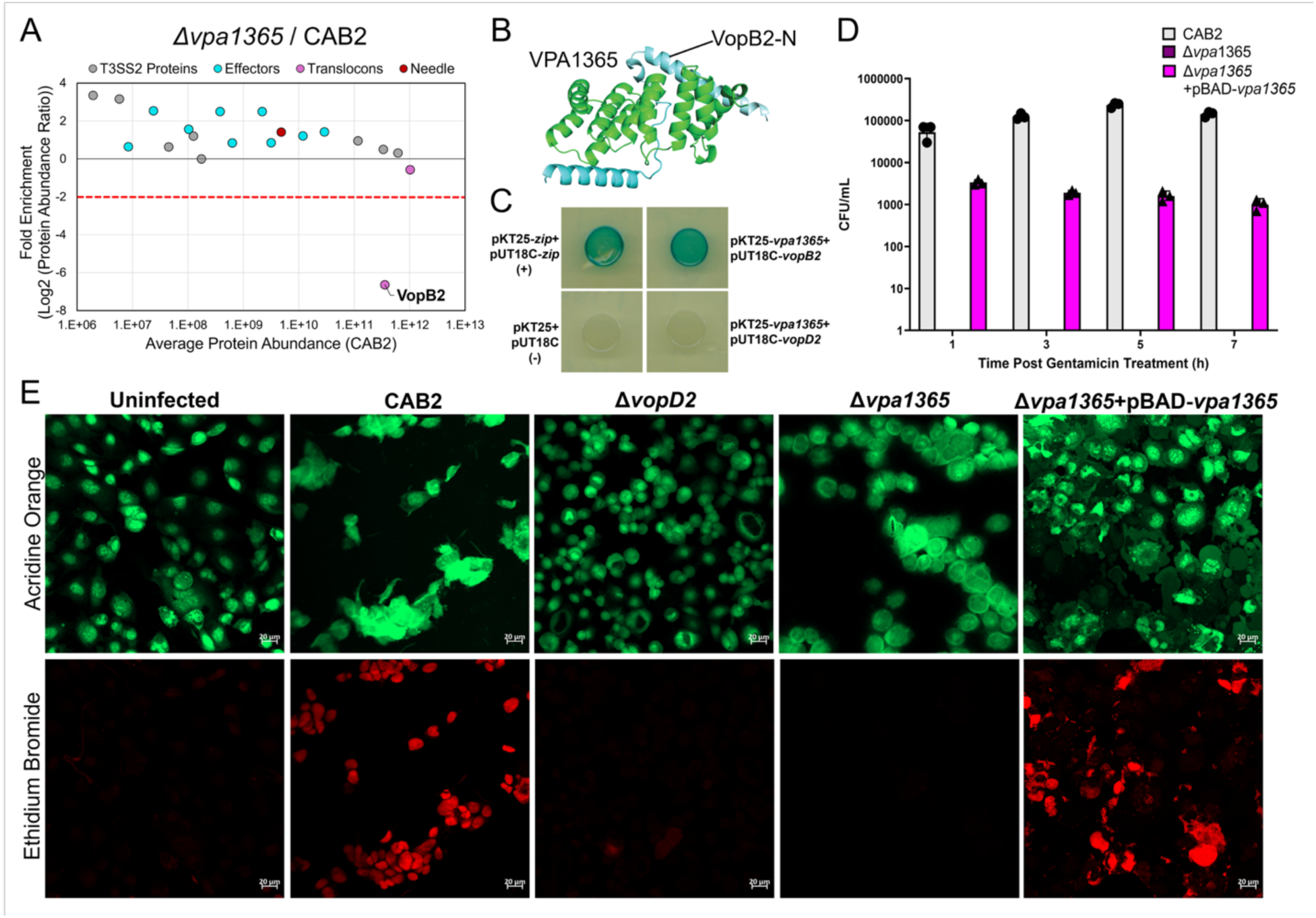
Absence of VPA1365 abolishes pore formation in Caco-2 cells. **(A)** Plot of T3SS2 proteins identified by label-free LC-MS/MS comparing Δ*vpa1365* and CAB2 secreted protein fractions. The fold enrichment (Δ*vpa1365*/CAB2) is plotted against the average protein abundance in the CAB2 strain (*n = 3*). T3SS2 effectors are highlighted in cyan, the needle protein in magenta, and the translocons in purple. The red horizontal dotted line indicates the significance threshold (Log_2_(Δ*vpa1365*/CAB2) < -2). **(B)** Structure showing the AlphaFold-predicted interaction between VPA1365 (green) and N-terminus of VopB2 (cyan) (ipTM=0.8). **(C)** Bacterial two-hybrid assay demonstrating the interaction between VPA1365 and VopB2. A positive interaction was observed, as indicated by the formation of blue colonies. No interaction was detected between VPA1365 and VopD2, as indicated by white colonies. The negative control (pKT25 + pUT18C) and positive control (pKT25-zip + pUT18C-zip) were included for comparison. **(D)** Intracellular survival of the indicated *V. para* strains in Caco-2 cells. Colony-forming units (CFU) were quantified following 1.5 h of infection, removal of extracellular bacteria by gentamicin treatment for 1.5 h, and enumeration at 1, 3, 5, and 7 h post-gentamicin treatment (PGT). **(E)** Representative fluorescence micrographs of uninfected or infected Caco-2 cells stained with acridine orange (green) and ethidium bromide (red). Cells with compromised membrane integrity are positive for ethidium bromide staining (red).

Having established that VPA1365 binds the translocon VopB2, we next examined the functional consequences of its deletion on cell invasion. Gentamicin protection assays revealed that *Δvpa1365* cells can no longer invade Caco-2 cells, with no observable colonies detected (**Fig 4D**). Complementation of the gene on a plasmid (Δ*vpa1365*+pBAD-*vpa1365*) can partially rescue the phenotype (**Fig 4D**). Since translocons form the pores that allow effectors to enter host cells, we tested the effect of *vpa1365* deletion on pore formation directly. It has been previously shown that deletion of either the major translocon *vopB2* or the minor translocon *vopD2* from *V. para* cells disrupts pore formation in Caco-2 cells (34). To test the effect of deletion of *vpa1365* on pore formation, Caco-2 cells were incubated with either *V. para* CAB2, translocon mutant (Δ*vopD2*), the chaperone deletion mutant (Δ*vpa1365*), or the complement (Δ*vpa1365*+pBAD-*vpa1365*) strain for 2 hours. Cells were then stained with a membrane-permeable dye (acridine orange-green) and a membrane-impermeable dye (ethidium bromide-red) to visualize membrane leakage (presence of red color). We observe a positive ethidium bromide staining only in cells incubated with *V. para* CAB2 or the complement Δ*vpa1365*+pBAD-*vpa1365* strains, indicating pore formation when VPA1365 is present (**Fig 4E**). In Δ*vpa1365* invaded cells, pore formation was not detected, suggesting that deletion of the translocon chaperone alone is sufficient to prevent formation of pores in host cells. Together, these results support that VPA1365 functions as a specific chaperone for the major translocon protein VopB2 and is essential to form pores in host cells, a critical step for secretion of effectors into host cells for bacterial invasion.

### VPA1340 governs the assembly of the T3SS2 apparatus

Our fourth candidate, VPA1340, is a 9 kDa protein encoded on an operon with two overlapping coding sequences of structural genes. To ensure our deletion strains did not affect the production of the structural proteins, we created a mutant strain *vpa1340*\* encoding a premature stop codon (**S4A Fig**). LC-MS/MS analysis of the secreted proteins from the mutant strain caused a drastic reduction in the abundance of most T3SS2 substrates as compared to the wild-type (**Fig 5A**). Western blotting confirmed that representative effectors VopL and VopA could not be detected in the secreted fractions from the *vpa1340\** mutant strain, while both effectors were detected in the cell pellet samples (**S4B Fig**). Complementation of *vpa1340* on a plasmid or chromosome restored the secretion of effectors (**S4B Fig**).

**Fig 5:**
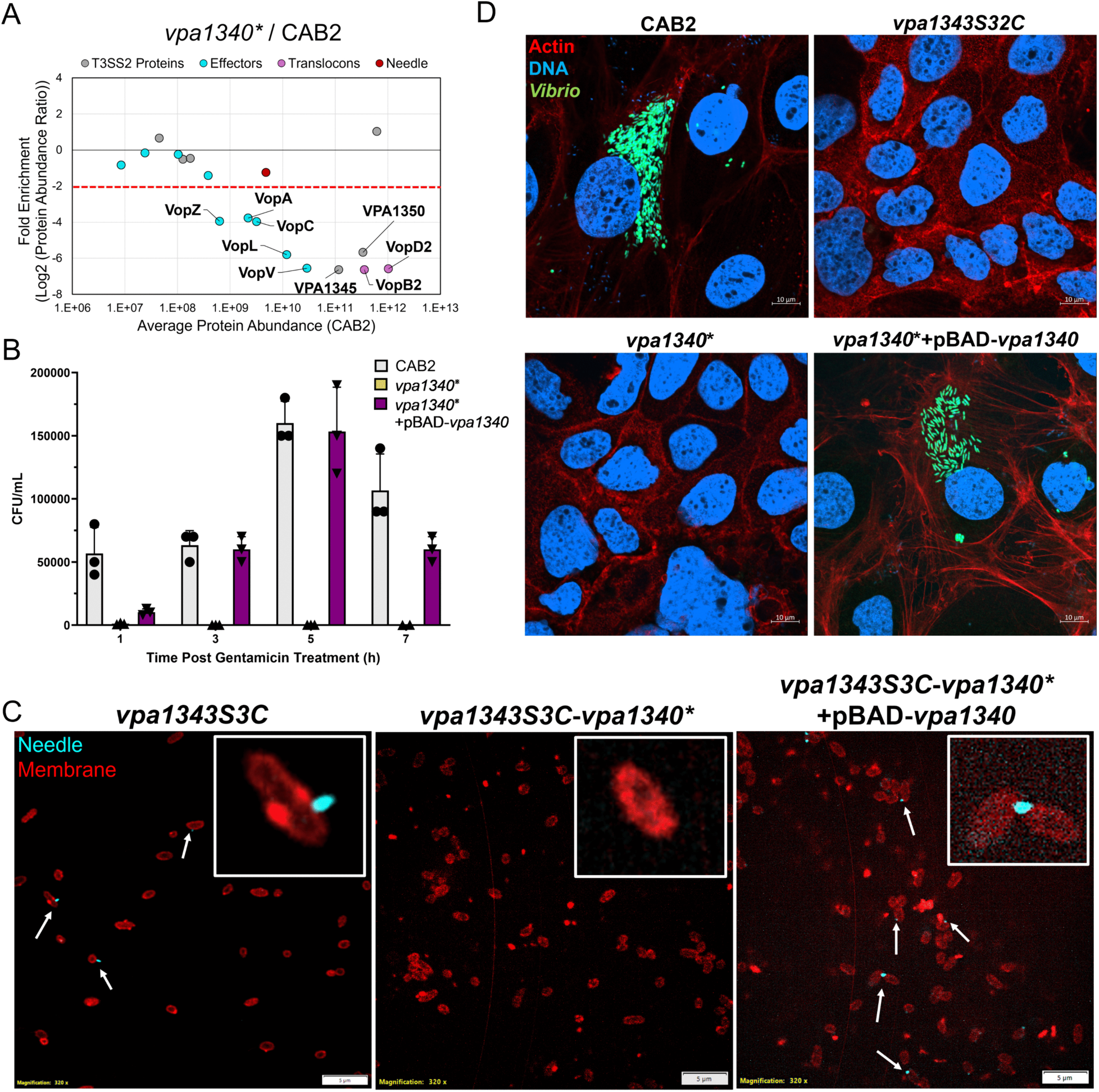
T3SS2 assembly is dependent on VPA1340. **(A)** Plot of T3SS2 proteins identified by label-free LC-MS/MS comparing *vpa1340*^⁎^ and CAB2 secreted protein fractions. The fold enrichment (*vpa1340*^⁎^/CAB2) is plotted against the average protein abundance in the CAB2 strain (*n = 3*). T3SS2 effectors are highlighted in cyan, the needle protein in magenta, and the translocons in purple. The red horizontal dotted line indicates the significance threshold (Log2(*vpa1340\**/CAB2) < -2). **(B)** Intracellular survival of the indicated *V. para* strains in Caco-2 cells. Colony-forming units (CFU) were quantified following 1.5 h of infection, removal of extracellular bacteria by gentamicin treatment for 1.5 h, and enumeration at 1, 3, 5, and 7 h post-gentamicin treatment (PGT). **(C)** Representative fluorescence micrographs showing T3SS2 needle structures (cyan) on the bacterial surface. Bacterial membranes were stained with Nile Red (red). Arrows indicate bacteria displaying detectable needle structures. **(D)** Representative confocal micrographs of Caco-2 cells at 5 h post-gentamicin treatment (PGT) infected with GFP-expressing intracellular *V. para* (green). Actin was stained with rhodamine-phalloidin (red) and DNA was stained with Hoechst (blue).

VPA1340 is a small protein with no defined domains that could provide insight into its function. The observed widespread effect of its disruption on effector secretion prevents prediction of potential interacting T3SS2 substrates. Additionally, Alphafold modeling of VPA1340 with each of the T3SS2 effectors did not yield strong interaction scores. Given that the reduction of T3SS2 effectors should severely impair bacterial invasion and survival in epithelial cells, we performed a gentamicin protection assay and confocal microscopy on Caco-2 cells infected with *V. para* CAB2, *vpa1340*\*, and the complement strain (*vpa1340*\*+pBAD-*vpa1340*). Bacteria with disrupted *vpa1340* were unable to invade Caco-2 cells, whereas complementation restored invasion and intracellular replication to levels similar to *V. para* CAB2 (**Fig 5B/ D**).

Collectively, these observations led us to hypothesize that VPA1340 may affect the assembly of the T3SS2 apparatus itself, thereby indirectly impacting effector secretion. To test this, we used a *V. para* mutant strain (*vpa1343S32C*), in which the needle protein, VPA1343, is mutated to prevent the secretion of effectors (35). LC-MS/MS analysis of the secreted proteins from this strain show that the secretion profile closely resembles that of the *vpa1340\** strain, in which most effectors are downregulated (**S5 Fig**). This result prompted us to investigate more directly whether VPA1340 affects needle formation in the bacteria. To visualize the needle, we utilized another previously characterized needle mutant strain (*vpa1343S3C*) that incorporates a cysteine into the needle to enable labeling with maleimide chemistry (35). High-resolution microscopy imaging showed that disruption of *vpa1340* completely impaired the formation of needles in *V. para* cells, while plasmid-based complementation of *vpa1340* restored needle structures (**Fig 5C**). For further validation, we used Western blot to analyze the presence of the needle protein in *vpa1340* mutants and as expected, the disruption of VPA1340 prevents production of VPA1343, while complementation restored it to normal levels (**S4C Fig**).

Absence of needle formation would affect the assembly of translocon proteins and thereby impair effector secretion, ultimately resulting in abortive infection. Consistent with impaired needle formation, confocal microscopy revealed that host cells infected with the non-secreting needle mutant (*vpa1343S32C*) displayed intact actin structures and no visible bacteria, indicative of failed infection similar to the *vpa1340\** mutant (**Fig 5D**). Together, these results indicate that VPA1340 is essential for T3SS2 needle assembly, and its disruption prevents effector secretion and host cell invasion.

## Discussion

Despite the central role of the T3SS2 in the enterotoxicity caused by *V. para*, the mechanisms governing its assembly and substrate hierarchy have remained unexplored. Here, we identify genes within the Vp-PAI7 pathogenicity island that function as stage-specific chaperones coordinating injectisome assembly and effector secretion. The chaperone-encoding genes are highly upregulated upon TDC induction. Using mass spectrometry analysis, our study reveals a hierarchical chaperone network that differentially regulates early (needle), middle (translocon) and late substrates (effectors) of T3SS2 and provides a mechanistic framework for its stepwise assembly.

A key feature of the T3SS is the ordered secretion of structural proteins and effectors, a process mediated by chaperones that bind substrates to stabilize them and facilitate secretion (2). While such chaperones have been well characterized in the T3SS1 of *V. para* (36, 37), their roles in T3SS2 have remained mostly unexplored. VPA1334 is the only reported T3SS2 chaperone to date, which was previously shown to function as a chaperone for the effector VopC (38). In this study, we combined reverse genetics with LC-MS/MS based methodology to identify substrates of T3SS2 chaperones. Specifically, we expand the chaperone repertoire of T3SS2 to include an additional substrate for the chaperone VPA1334 as well as three new chaperones: VPA1352, VPA1365, and VPA1340. Our data demonstrates that these chaperones regulate distinct stages of injectisome assembly and effector secretion. Precisely, we show that VPA1340 is required for assembly of the needle filament protein VPA1343 during early substrate secretion, forming the conduit for effector translocation. VPA1365 functions at the next stage by facilitating assembly of the major translocon protein VopB2, enabling host cell contact and pore formation. Finally, VPA1334 and VPA1352 act as chaperones for effectors VopC, VopL and VopV, all of which are critical for the pathogenicity of the bacterium (**Fig 6**).

**Fig 6:**
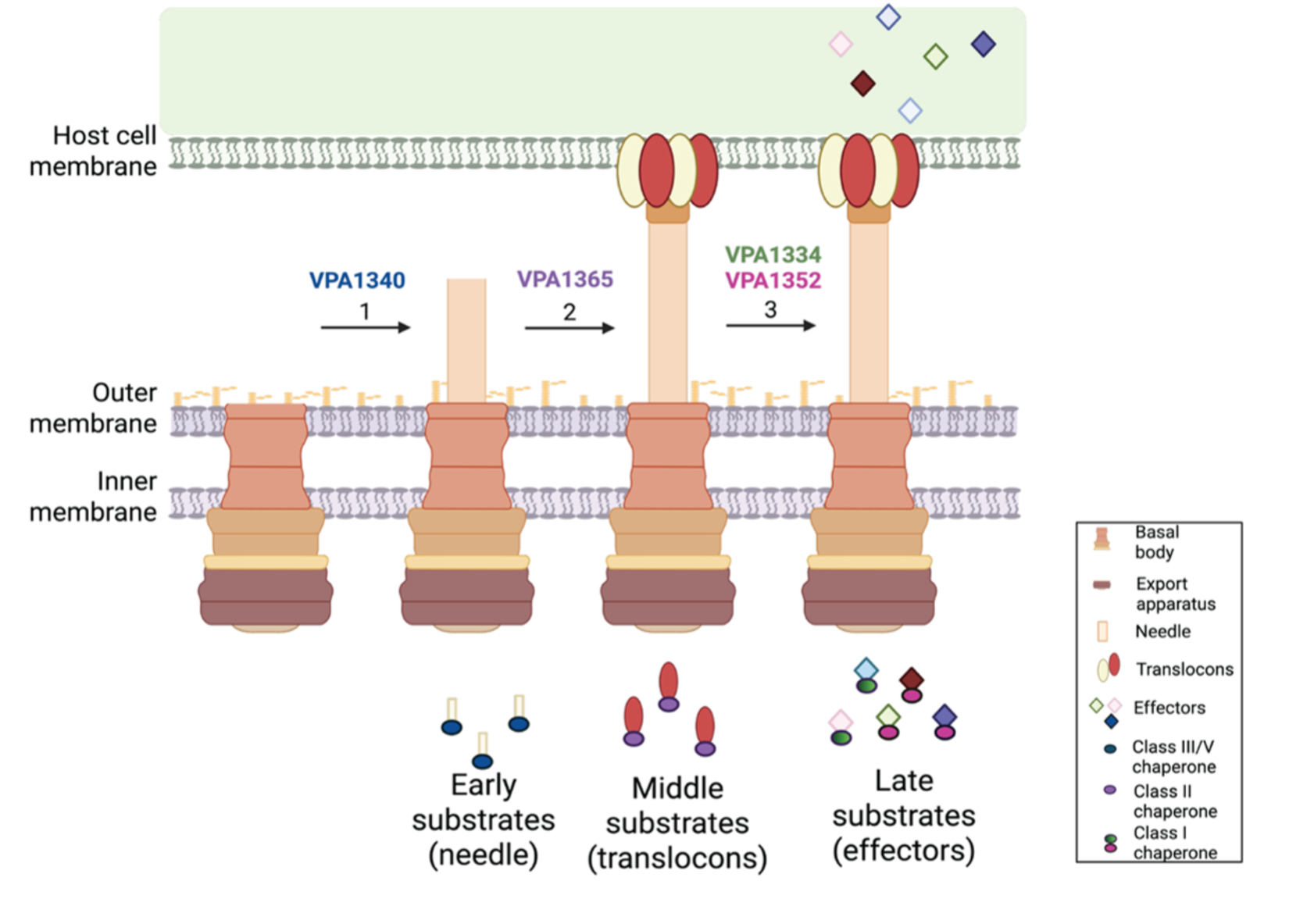
Schematic representation of T3SS2 chaperones in injectisome assembly and substrate secretion. The T3SS2 spans the bacterial inner and outer membranes and extends outward to contact the host cell membrane, forming a conduit for protein translocation. Substrates are secreted in a defined hierarchy of early, middle, and late substrates. Early substrates are responsible for assembly of the needle structure. Middle substrates mediate formation of the translocon pore in the host cell membrane. Late substrates consist of effectors that are delivered into the host cytoplasm. The *V. para* T3SS2 chaperones (VPA1340, VPA1365, VPA1334 and VPA1352) are indicated at the stages of T3SS2 assembly where they function. Created in BioRender. Casey, A. (2026) https://BioRender.com/tk13o7m

The effector chaperones VPA1334 and VPA1352 have a similar fold with three-α helices and five-β-strands, consistent with their function as class I effector chaperones (**S2 Table**) (39). The available information about class 1A effector chaperones suggests that they are located next to each of the effectors that they regulate, similar to that seen for *Salmonella typhimurium* (33), *Yersinia enterocolitica* (40), enteropathogenic *E. coli* (41) and the T3SS1 of *V. para* (36). However, we show that VPA1334 (likely a class 1B chaperone) is located in the pathogenicity island but distantly located from the effectors (VopC and VopL) it regulates. Deletion of *vpa1334* leads to a significant decrease in the secreted levels of effectors VopC and VopL in induced bacterial cells, suggesting that binding of VPA1334 leads to the stabilization of these effectors in the cytoplasm prior to secretion. The second effector chaperone VPA1352 is in close proximity, but not directly adjacent to its effector VopV. Deletion of *vpa1352* affects the secretion of VopV drastically; however, VopV levels are not significantly affected in the cell pellet of the Δ*vpa1352* strain. This result mirrors results for the *Yersinia enterocolitica* T3SS, where deletion of the SycH chaperone leads to accumulation of effector YopH in the cytoplasm but a reduction of YopH secretion (42).

Chaperones VPA1334 and VPA1352 facilitate secretion of effectors VopC, VopL and VopV, which collectively target the host actin cytoskeleton. VopC deamidates Rac1 to facilitate membrane ruffling and invasion of *V. para* into epithelial cells (22) while VopL induces the formation of stable actin filaments (43). In the absence of VopL, bacteria undergo filamentation inside epithelial cells (23). Concurrently, we observe that in the absence of its cognate chaperone VPA1334, bacterial filamentation is also observed in infected Caco-2 cells. The F-actin binding effector VopV mediates actin bundling and is proposed to be responsible for T3SS2-dependent enterotoxicity in a rabbit ileal loop model (21). However, its contribution to invasion in epithelial cells was not reported. We show deletion of *vopV* or its chaperone *vpa1352* does not significantly affect the invasion and replication of bacteria in Caco-2 cells. Collectively, our findings demonstrate that the chaperones VPA1334 and VPA1352 regulate three effectors that specifically target the actin cytoskeleton. The activities mediated by VopL and VopC, but not VopV, are required for efficient invasion and dissemination, thereby positioning these effectors among non-redundant core components of T3SS2-mediated virulence. We demonstrate VopC, VopL, and VopV effectors for *V. para* are coupled to dedicated chaperones that ensure their access to the T3SS2 secretion machinery.

VPA1365 is a 19 kDa protein and a member of the TPR family that is annotated as a putative two-component response regulator. The basic function of TPR folds is to mediate protein-protein interactions (44), and both needle and translocation chaperones adopt this fold (13–15). However, a recent study by Yin et al. reported that VPA1365 binds to the promoter of *vtrA* and increases the expression of T3SS2 genes. They also report that VPA1365 binds to the promoter of many additional virulence-associated genes such as *scrG*, *pilA*, and *mshA* (45). However, our study reveals an alternative function for VPA1365 as a class II chaperone within T3SS2. Unlike its previously described role in transcription regulation, we demonstrate that VPA1365 is essential for the assembly of the major translocon protein VopB2. This result is supported by the fact that deletion of *vpa1365* specifically disrupts the secretion of VopB2 without affecting the secretion of any other proteins encoded in Vp-PAI7. Further, our data show that absence of VPA1365 prevents pore formation in Caco-2 cells without affecting the secretion of effectors thus validating the function of VPA1365 as a class II chaperone (**S2 Table**).

The open reading frame of the gene *vpa1340* encodes a 9 kDa protein containing two predicted *a*-helices. Due to its small size and low-confidence structure prediction, its function has remained elusive. The gene is encoded on an operon between two T3SS2 structural proteins. We show that disruption of *vpa1340* by introducing a premature stop codon disrupts secretion of most of the T3SS2 substrates. The secretion of all T3SS2 substrates was significantly downregulated in the *vpa1340\** strain, impairing bacterial invasion and replication in epithelial cells. This phenotype closely resembles that of the T3SS2 needle mutation, where a single S32C mutation in the needle protein VPA1343 impairs the secretion of many T3SS2 substrates(35). Our results show that VPA1340 plays a critical role in stabilizing the needle protein in the cell, as the disruption of *vpa1340* results in complete loss of the needle protein in the cytoplasm. Interestingly, we initially expected that VPA1340 may be interacting directly with the needle protein VPA1343 to form a complex, like the other chaperone behaviors reported here. However, when we tested this potential using Alphafold modeling and the BACTH assay, it did not appear that VPA1340 and VPA1343 can form a stable complex. It has been reported in other bacteria that needle formation is dependent on binding from two chaperone proteins, one of which resembles a small helical protein similar to VPA1340 (14, 46, 47). This potential co-chaperone complex could explain the absence of a direct interaction between VPA1340 and VPA1343. Collectively our results indicate that VPA1340 is required for the stabilization of the needle protein VPA1343, functionally grouping VPA1340 as a class V chaperone involved in needle biogenesis (**S2 Table**).

Previous studies have largely characterized T3SS chaperones as substrate-specific factors that stabilize cognate proteins and facilitate their delivery to the secretion apparatus (11, 48–52). The four T3SS2 chaperones identified in *V. para* represent the major functional classes of T3SS chaperones. VPA1334 and VPA1352 are associated with effectors and their folds are consistent with class I chaperones, which commonly stabilize effectors in the bacterial cytoplasm and facilitate their targeting to the secretion apparatus (53). VPA1365 is associated with a translocon protein and functions as a class II chaperone. Such chaperones are important for maintaining the hydrophobic translocon substrates in a secretion-competent state and contribute to preferential secretion of translocon proteins before effectors (49, 54). Finally VPA1340 is linked to the needle protein, suggesting a class V function in preventing premature needle-subunit assembly and supporting early injectisome biogenesis (13).

Disruption of the needle associated chaperone VPA1340 broadly impairs protein secretion, establishing it at the forefront of the chaperone hierarchy. Next, VPA1365 acts as a sequential checkpoint that regulates pore formation in host cells during invasion. Ultimately, VPA1334 and VPA1352 define parallel pathways for substrate delivery. Specifically, our findings indicate that effectors VopC, VopL, and VopV each have cognate chaperones for secretion. Our systematic analysis of all three chaperone classes reveals coordinated but distinct contributions to successive stages of injectisome assembly. The differential defects observed following deletion or disruption of individual chaperone classes support a model in which hierarchical secretion is regulated. The presence of these substrate-associated chaperones in *V. para* is consistent with the hiearchial organization of T3SS secretion, in which needle components are secreted first, followed by translocon proteins and finally effectors (55). These findings contribute to the growing body of knowledge surrounding chaperone proteins across various pathogenic bacteria (**S2 Table**). Together, these findings support a model in which specific chaperone-substrate interactions contribute to the organization and hierarchy of T3SS substrate deployment. This further highlights the potential of *V. para* as a model system for studying T3SS chaperones and identifying conserved chaperone-mediated mechanisms in other bacteria.

T3SS2 has been long recognized as essential for both invasion and enterotoxicity in *V. para*. The elucidation of the roles of previously uncharacterized proteins that act as regulators of the T3SS2 secretion hierarchy expands the regulatory repertoire of T3SS2. The presence of multiple chaperones within the T3SS2 reflects the complex challenges faced by the secretion system, which must coordinate different stages of assembly and infection. No single chaperone can carry out all aspects of this process, necessitating the employment of a hierarchical network. Our findings reveal new nodes within this network that may serve as potential targets for disrupting or inhibiting T3SS2 function during infection. Since the activity of T3SS2 directly corresponds to the infection phenotype, the potential of targeting T3SS2 chaperones presents an attractive therapeutic strategy. Future studies will aim to uncover the molecular mechanisms of the chaperone-substrate interactions through structure-guided methods. Ultimately, the findings presented in this study provide a framework for identifying new ways in which secretion can be disrupted to control the T3SS2-mediated bacterial pathogenicity.

## Materials and Methods

### Bacterial strains and growth conditions

The *Vibrio parahaemolyticus (V. para)* CAB2 strain was derived from the POR1 strain (clinical isolate RIMD2210633 lacking TDH toxins, and was generously gifted by Drs. Tetsuya Iida and Takeshi Honda (56). *E. coli* Mach1 was used for transformation of pBAD plasmids and strain S17 was used for pDM4 plasmids. All *E. coli* strains were cultivated in Lysogenic Broth (LB) at 37℃ while *V. para* strains were grown in MLB (LB medium supplemented with 3% NaCl (w/v)) at 30℃ for routine cultivation. When necessary, antibiotics were added to maintain plasmids-ampicillin 100 µg/ml, kanamycin 30 µg/ml, chloramphenicol 25 µg/ml for *E. coli* and spectinomycin 50 µg/ml and kanamycin 250 µg/ml for *V. para*.

### Plasmid and strain construction

For in-frame deletion or modification of genes in the chromosome of *V. para*, allelic exchange using suicide vector, pDM4 was employed. For deletion of *vpa1334*, primer pairs JJ237/236 and JJ235/240 were used to amplify the upstream and downstream flanking regions of *vpa1334*, fragments joined by overlap PCR (JJ237/240), digested with XbaI/SalI and ligated to pDM4 digested with the same enzymes. To delete *vpa1352*, primer pairs JJ387/388 and JJ389/390 were used to amplify the upstream and downstream flanking regions of *vpa1352*, fragments joined by overlap PCR (JJ387/390), digested with NheI/XhoI and ligated to pDM4 digested with the same enzymes. To delete *vpa1365*, primer pairs JJ343/345 and JJ342/344 were used to amplify the upstream and downstream flanking regions of *vpa1365*, fragments joined by overlap PCR (JJ342/345), digested with NheI/XhoI and ligated to pDM4 digested with the same enzymes. To disrupt *vpa1340*, primer pairs JJ209/296, JJ297/298, JJ299/212 and were used to amplify the upstream, mutated coding sequence and downstream regions of *vpa1340*, fragments joined by overlap PCR (JJ209/212), digested with NheI/XhoI and ligated to pDM4 digested with the same enzymes. The sequences of the primers are listed in **S3 Table**. The cloned products were transformed to *E. coli* S17. After successful transformation, the *E. coli* S17 strain containing the plasmid was conjugated to *V. para* CAB2 strain and the transconjugants were selected on minimal marine medium (MMM) agar containing 25 µg/ml chloramphenicol. For curing *sacB*-containing pDM4, the conjugants were counter-selected on MMM agar containing 15% (w/v) sucrose. Successful deletion or modification was confirmed by PCR and sequencing.

For complementation, the coding sequence of each gene was amplified using primer pairs JJ453/454 and JJ455/456 for *vpa1334* and *vpa1352*, respectively, digested with BglII/HindIII and ligated into pBAD-Fix digested with the same enzymes. The coding sequence of *vpa1365* was amplified using primer pairs JJ360/361, digested with Xho1/KpnI and ligated into pBAD-Fix digested with the same enzymes. The coding sequence of *vpa1340* was amplified using primer pairs JJ230/231, digested with Xho1/EcoRI and ligated into pBAD-Fix digested with the same enzymes. The resultant pBAD-Fix constructs were transformed to *E. coli* Mach1 and introduced to the *Vibrio* strain via triparental conjugation using *E. coli* DH5ɑ containing pRK2073 (Kan^R^). The transconjugants were selected on MMM agar containing 250 µg/ml kanamycin. Expression of the genes was induced by adding 0.02% arabinose. The plasmids and strains constructed in the study are listed in **S4 Table**.

### RNA extraction and sequencing

Overnight cultures of CAB2 were sub-cultured to an OD600 of 0.3 in MLB media. Three biological replicates of CAB2 under uninduced conditions and with 100 µM taurodeoxycholate (TDC) induction were grown for 1.5 hours at 37℃. Samples were normalized to an OD600 of 1.0 and RNA extraction was performed using a RNeasy Mini kit (Qiagen) according to manufacturer’s instructions, with minor modifications. Briefly, cells were harvested and resuspended in TE buffer containing 1 mg/ml lysozyme. After incubation for 10 minutes at room temperature, 700 µl of buffer RLT was added to each tube and vortexed vigorously, followed by addition of 500 µl of ethanol. The mixture was transferred to a RNeasy Mini spin column, then centrifuged, washed and eluted according to the protocol. Following NanoDrop quantification, DNase treatment was performed as per manufacturer’s instructions (Thermo Fischer Scientific, USA). Samples were shipped on dry ice, and and reads were generated at Azenta Life Sciences using Illumina 2x150bp whole transcriptome sequencing via rRNA depletion. Sequenced reads were trimmed using Trimmomatic v.0.36 and mapped to the reference *Vibrio parahaemolyticus* RIMD2210633 genome (GCA_000196095_1) using the Bowtie2 aligner v.2.2.6. Unique gene hit counts (only within the gene regions) were calculated using featureCounts from the Subread package v.1.5.2. Hit counts were summarized and reported using gene_id feature of the annotation file. Gene hit counts (filtered at 1) for the three TDC samples were compared to those of the three uninduced samples using DEseq2 v.1.38.3.

### Secretion Assay

Overnight cultures of bacteria were sub-cultured to an OD600 of 0.3 in LB media supplemented with 20 mM KCl. 100 µM taurodeoxycholate (TDC) was added to induce T3SS2 and the bacteria were grown at 37℃ for 3 hours. When necessary, arabinose was added at a concentration of 0.02%. After 3 hours, to collect the pellet fraction, bacterial cultures of OD600 of 0.5 were pelleted and resuspended in 2X Laemmli buffer. For the secretion fraction, the cultures were normalized to the lowest OD among samples and spun at 4000 rpm, 15 min at 4℃. The supernatant was filtered through a 0.22 µm filter into chilled conical tubes. Deoxycholate (150 µg/ml), and Trichloroacetic acid (7% v/v) were added to the supernatant, mixed by inverting, and protein precipitation was allowed to proceed at 4℃ overnight. The following day, the precipitated proteins were collected by spinning the tubes at 14000 rpm for 15 min at 4℃, and the pellet was washed twice with ice-cold acetone. After air-drying for 30min, the pellet was resuspended in 2X Laemmli buffer.

### Liquid Chromatography-Tandem Mass Spectrometry Analysis

Protein samples from the following *V. para* strains were analyzed by mass spectrometry (n = 3 biological replicates for each): CAB2, Δ*vpa1365*, *vpa1340*\*, Δ*vpa1334*, Δ*vpa1352*, Δ*vpa1334*Δ*vpa1352* and *vpa1343S32C*. Pellet and secreted protein fractions from each strain were resolved by SDS-PAGE gels and stained with Coomassie blue dye. Samples were reduced with 10 mM DTT for 1 hour at 56°C and then alkylated with 50 mM iodoacetamide for 45 min at room temperature in the dark. Trypsin digestions were performed overnight at 37°C (MS-grade, Thermo Scientific). Resulting peptide samples were de-salted via solid phase extraction (SPE) prior to LC-MS/MS analysis. Experiments were carried out on a Thermo Scientific EASY-nLC coupled to an Orbitrap Fusion Lumos mass spectrometer (Thermo Fisher Scientific). Peptide samples were separated on a 50 cm Thermo PepMap analytical column (2 µm particle size, 75 µm diameter) (Thermo Fisher Scientific) over a 100 min gradient. MS1 spectra were acquired in the Orbitrap mass analyzer (resolution 120,000) and peptide precursor ions were isolated and fragmented using high-energy collision-induced dissociation (HCD) to generate MS/MS spectra which were collected in the ion trap.

Data was searched using Proteome Discoverer version 2.2 software (Thermo) against sequences in the *Vibrio parahaemolyticus* serotype O3:K6 (strain RIMD 2210633) protein database (Taxon ID: 223926). Seach parameters included the following: precursor mass tolerance of 15 ppm, fragment ion mass tolerance of 0.6 Da, 2 missed cleavages allowed, variable modifications of Carbamidomethylation (cysteine) (+57.021 Da), oxidation (methionine) (+15.995 Da), and acetylation (protein N-termini) (+42.011 Da). Proteins were filtered to a false discovery rate (FDR) threshold of 1% at both the peptide spectral match level and protein level. Additionally, all proteins must have at least two unique peptide sequence identifications to be included. Protein abundances were calculated by label-free quantitation using precursor ion area values. Protein abundance ratios were calculated using a summed abundance approach from 3 biological replicates of each type. Missing abundances were replaced with random values sampled from the lower 5% of detected values. P-values were calculated using an ANOVA test based on individual proteins (PD 2.2). Significantly enriched proteins (shown in red in volcano plots) have a Fold Change > 1.5 and p-value < 0.05, while significantly decreased proteins (shown in blue in volcano plots) have a Fold Change < 0.5 and p-value < 0.05.

### Protein-protein interactions

To visualize chaperone interactions with their preferred substrates, we submitted the corresponding sequences to the AlphaFold3 server. We modeled two chains for VPA1334 and VPA1352 interactions with their corresponding effectors to identify potential translocation signal sequences for the effectors. We then generated AlphaFold interaction models for the trimmed effector translocation sequences to report confidence metrics (ipTM). To experimentally characterize the interaction between the chaperones (VPA1334, VPA1352 and VPA1365) with their substrates, Bacterial Adenylate Cyclase-based Two-Hybrid System (BACTH, Euromedex) was employed. Primer pair JJ435/436 was used to amplify *vpa1334*, digested with KpnI/EcoR1 and ligated to pKT25 digested with the same enzymes. Primer pair JJ457/458 was used to amplify *vpa1352*, digested with KpnI/EcoR1 and ligated to pKT25 digested with the same enzymes. Primer pair JJ489/490 was used to amplify *vpa1365*, digested with XbaI/KpnI were ligated into pKT25 digested with the same enzymes. Coding sequence corresponding to 1-200 amino acids of *vopV* was amplified using JJ459/460, digested with XbaI/EcoRI and ligated into pUT18C digested with the same enzymes. Primer pair JJ476/478 was used to amplify *vopC*, digested with BamHI/EcoRI and ligated into pUT18C digested with the same enzymes. Primer pair JJ439/440 was used to amplify *vopL*, digested with XbaI/EcoRI and ligated into pUT18C digested with the same enzymes. Primer pair JJ491/492 was used to amplify *vopD2*, digested with XbaI/EcoRI and ligated into pUT18C digested with the same enzymes. Primer pair JJ493/494 was used to amplify *vopB2*, digested with XbaI/EcoRI and ligated into pUT18C digested with the same enzymes. Sequences of primer pairs are mentioned in **S3 Table**. *E. coli* BTH101 was co-transformed with pKT25 and pUT18C plasmids containing the gene of interest and transformants were selected on LB agar plates supplemented with 40 μg/ml 5-bromo-4-chloro-3-indoyl-β-D-galactopyranoside (X-Gal), 0.5 mM isopropyl-β-D-thiogalactopyranoside (IPTG), 100 μg/ml ampicillin and 30 μg/ml kanamycin. Positive interactions were detected as blue colored colonies after incubation for 3-4 days at 30°C.

### Cell culture, gentamicin protection assay and microscopy

Caco-2 cells (ATCC) were cultured in MEM/EBSS with 20% Fetal bovine serum (FBS), 1% Penicillin and Streptomycin (PenStrep) for routine culture. Gentamicin protection assays were performed as described previously (57). Briefly, *V. para* strains were induced with TDC for 1.5 hours and then allowed to infect Caco-2 cells for 1.5 hours prior to gentamicin treatment to kill all extracellular *V. para*. Cells were then lysed at 1, 3, 5, or 7 hours post-gentamicin treatment (PGT) to quantify intracellular bacteria. For microscopy experiments, the wild-type and mutant strains were transformed with plasmid pMW-GFP, constitutive GFP expression plasmid. Caco-2 cells were seeded at a density of 2*10^5^ cells/ml on sterile poly-L-Lysine coated coverslips (Neuvitro Corporation, USA) and allowed to adhere for 48 hours. These cells were infected with the indicated *V. para* strains and treated with gentamicin as described above. Cells from the 5 hours PGT time point were washed twice with PBS and fixed with 3.2% (v/v) paraformaldehyde for 10 min at room temperature with gentle shaking. Cells were then washed once with PBS and permeabilized using 0.1% Triton-X-100 for 10 min. After washing with PBS, the coverslips were mounted on to glass slides with ProLong Gold Antifade Mountant and cured overnight. All imaging was performed on a Zeiss LSM 710 confocal microscope.

### Western Blotting

Following the secretion assay, equal volumes of pellet and supernatant samples were suspended in 5X SDS sample buffer and resolved on Tris-Glycine gels with the BioRad Mini-PROTEAN Tetra Cell system. Proteins were transferred onto Millipore Immobilon-P 0.45μM pore size polyvinylidene difluoride membrane using an Invitrogen Power blotter Semi-Dry transfer system. Membranes were rewetted with 100% methanol and rocked in blocking buffer (5% nonfat dry milk) 1X TTBS overnight at 4°C. Primary antibodies (rabbit anti-VopL (1:3000), rabbit anti-VopA (1:1000), and rabbit anti-VPA1343 (1:1000)) were diluted in the blocking buffer and added to the membranes with gentle shaking for 1 hour at room temperature. Membranes were then washed in 1X TTBS three times. Secondary antibody (donkey anti rabbit HRP conjugated (1:10,000)) was diluted in blocking buffer and added to membranes with gentle shaking for 1 hour at room temperature. Membranes were then washed in 1X TTBS three times and developed using Pierce ECL Western Blotting substrate (Thermo Scientific) and visualized using ChemiDoc Imaging System (Bio-Rad).

### Pore formation assay

Caco-2 cells were seeded at a density of 2*10^5^ cells/ml on sterile poly-L-Lysine coated coverslips (Neuvitro Corporation, USA) and allowed to adhere for 48 hours. *V. para* strains were induced with TDC for 1.5 hours and subsequently used to infect the Caco-2 cells at an MOI of 100. After 2 hours of infection, the cells were washed twice with PBS, followed by the addition of 1ml PBS containing 25 μg/ml ethidium bromide and 5 μg/ml acridine orange. The cells were incubated for an additional 15 minutes. After washing with PBS, the coverslips were mounted on to glass slides with ProLong Gold Antifade Mountant and allowed to cure overnight. The following day, the cells were imaged using a Zeiss LSM 710 confocal microscope.

### Microscopy of bacterial needle structures

A thiol chemistry-based approach was used to tag and visualize the formation of T3SS2 needles, as described previously (35). *V. para* strains were incubated with 100µM TDC for 25 minutes at 37 ℃. Nile red (1 µg/ ml) and Alexa Fluor 647 C_2_ Maleimide (10 µM) were added to the cells and incubated further for 20 minutes at 37℃. The bacterial cells were harvested by centrifugation at 5000 rpm for 5 minutes, and the cells were fixed with 3.2% paraformaldehyde. After fixation, the cells were washed twice with PBS and spotted on poly-L-Lysine coated coverslips. The coverslips were mounted on to glass slides with ProLong Gold Antifade Mountant and allowed to cure overnight. Images were acquired the next day using the Olympus Spin-SR Spinning Disk Confocal Microscope System at x320 magnification.

## Acknowledgements

We thank the members of the Orth lab for their discussions and editing. We also thank Euromedex and Dr. Zezhang T. Wen, Louisiana State University Health Sciences Center for providing the Bacterial-two hybrid system plasmids.

