## Supplementary material for "Distinct chaperones coordinate the type III secretion system assembly and effector secretion in *Vibrio parahaemolyticus*": supp file

### Supporting information

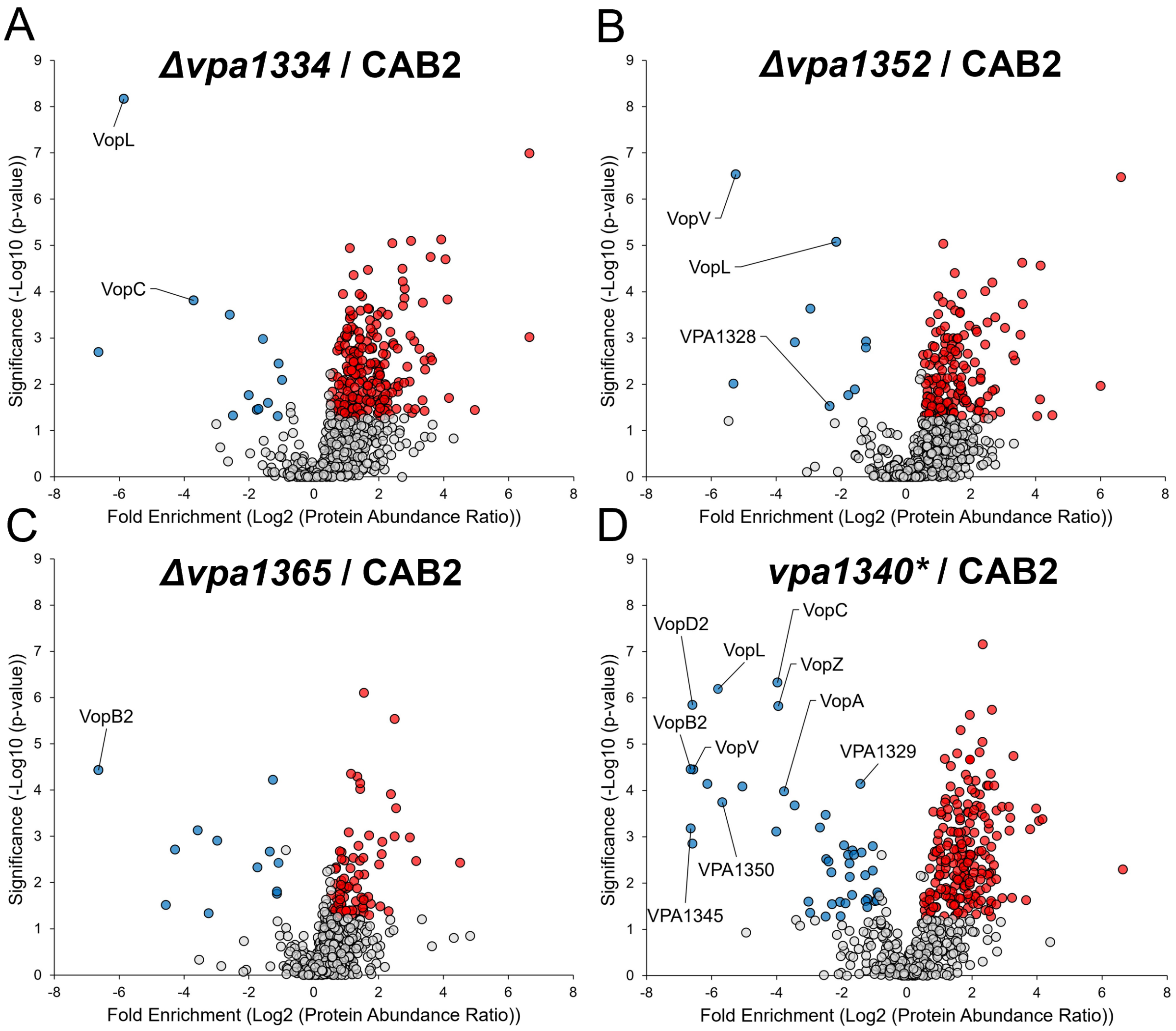

**S1 Fig. Effect of candidate chaperone gene deletion on *V. para* protein secretion.**

Volcano plots depicting the fold enrichment ($Log_2$(protein abundance ratio) of (mutant / wild-type) versus significance ($-Log_{10}$(p-value)) of secreted proteins identified by label-free LC-MS/MS from **(A)** Δ*vpa1334,* **(B)** Δ*vpa1352,* **(C)** Δ*vpa1365,* and **(D)** *vpa1340** compared with CAB2 following TDC induction. Proteins significantly enriched in the mutant strains are highlighted in red and defined by a fold change > 1.5 with a p-value < 0.05, while proteins significantly depleted are highlighted in blue and defined by a fold change < 0.5 with a p-value < 0.05 (*n = 3*). Significantly depleted T3SS2 proteins are labelled in each volcano plot.

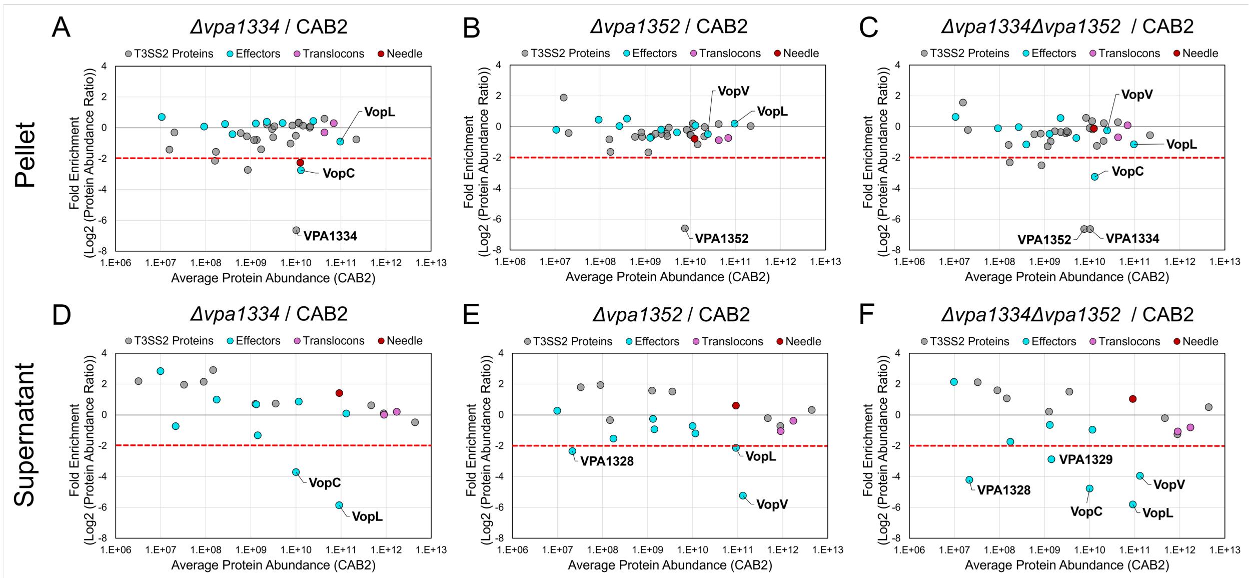

**S2 Fig. T3SS2 proteins identified by label-free LC-MS/MS data in pellet and supernatant fractions.**

Comparison of LC-MS/MS data of pellet and supernatant fractions from Δ*vpa1334*, Δ*vpa1352*, and Δ*vpa1334*Δ*vpa1352* strains. Pellet fractions: (**A**) Δ*vpa1334* vs. CAB2, (**B**) Δ*vpa1352* vs. CAB2 and (**C**) Δ*vpa1334*Δ*vpa1352* vs. CAB2. Supernatant fractions: (**D**) Δ*vpa1334* vs. CAB2, (**E**) Δ*vpa1352* vs. CAB2 and (**F**) Δ*vpa1334*Δ*vpa1352* vs. CAB2 strains. Fold enrichment (Log2(Protein Abundance Ratio) of mutant / CAB2 is plotted against the average protein abundance in the CAB2 strain (*n = 3)*. T3SS2 effectors are highlighted in cyan, the needle protein in magenta, and the translocon protein in purple.

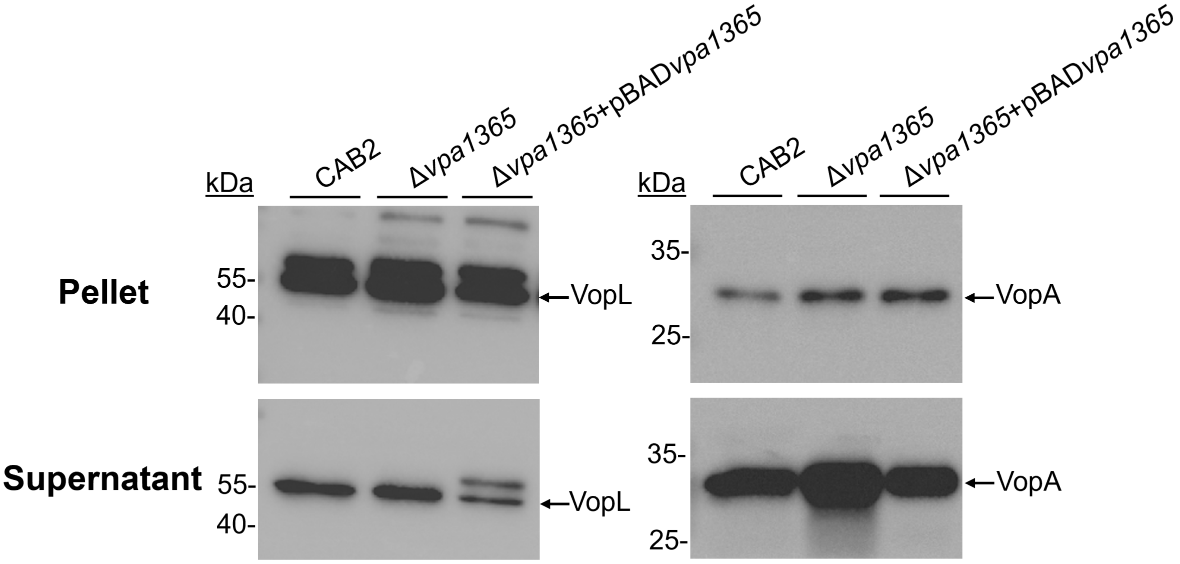

**S3 Fig. Western blot analysis of the T3SS2 effectors VopL and VopA in Δ*vpa1365* strains.**

Western blot of T3SS2 effectors VopL and VopA in the CAB2, Δ*vpa1365* and Δ*vpa1365*+pBAD-*vpa1365* strains grown under TDC induced condition followed by staining with anti-VopA or anti-VopL antibodies. Both VopL and VopA are observed in the supernatant fraction indicating secretion of these effectors.

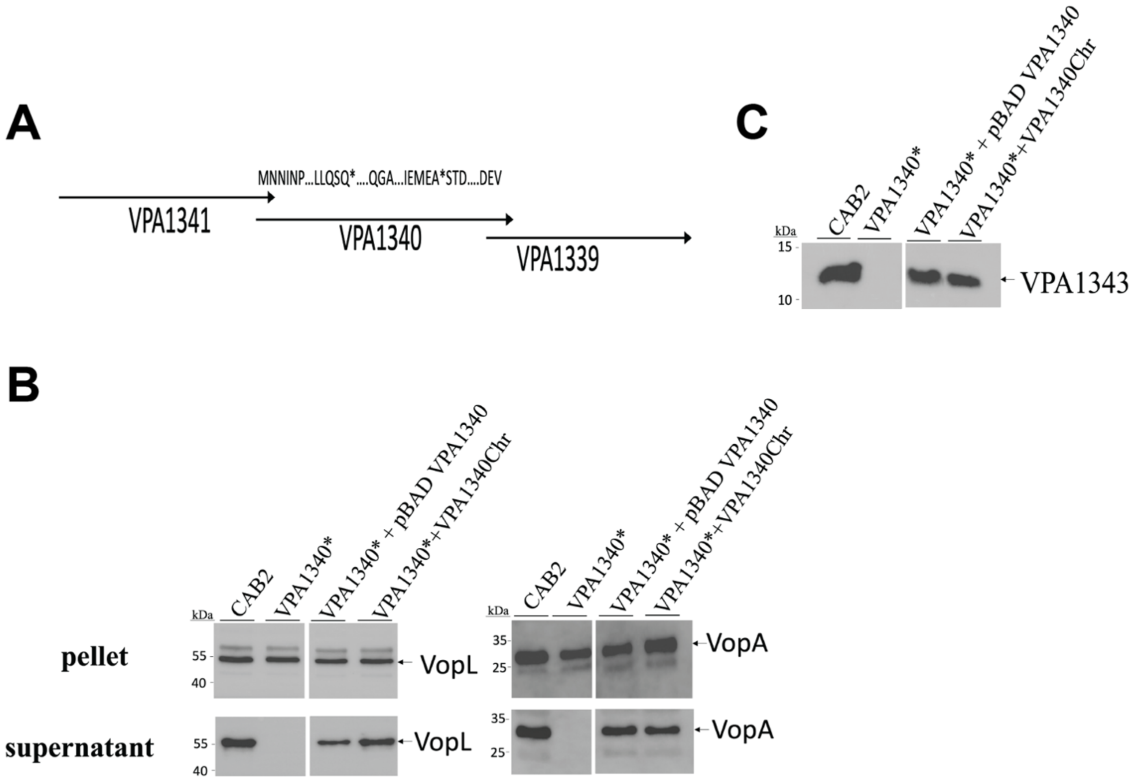

**S4 Fig. Disruption of *vpa1340* alters T3SS2 needle protein levels in the pellet fraction without affecting the expression of effectors.**

**(A)** Schematic illustrating the strategy used to generate the in-frame deletion of *vpa1340*. The deleted region and genomic organization are indicated. **(B)** Western blot analysis of pellet and supernatant from CAB2, *vpa1340**, *vpa1340**+pBAD- *vpa1340* and *vpa1340**+*vpa1340chr* strains showing that disruption of *vpa1340* does not alter the expression of the indicated T3SS effectors but specifically impairs the secretion. **(C)** Western blot analysis of the T3SS2 needle protein VPA1343 showing altered protein levels in the pellet fraction of *vpa1340** mutant compared with CAB2 and complement strains *vpa1340**+pBAD- *vpa1340* and CAB2*vpa1340**+*vpa1340chr*.

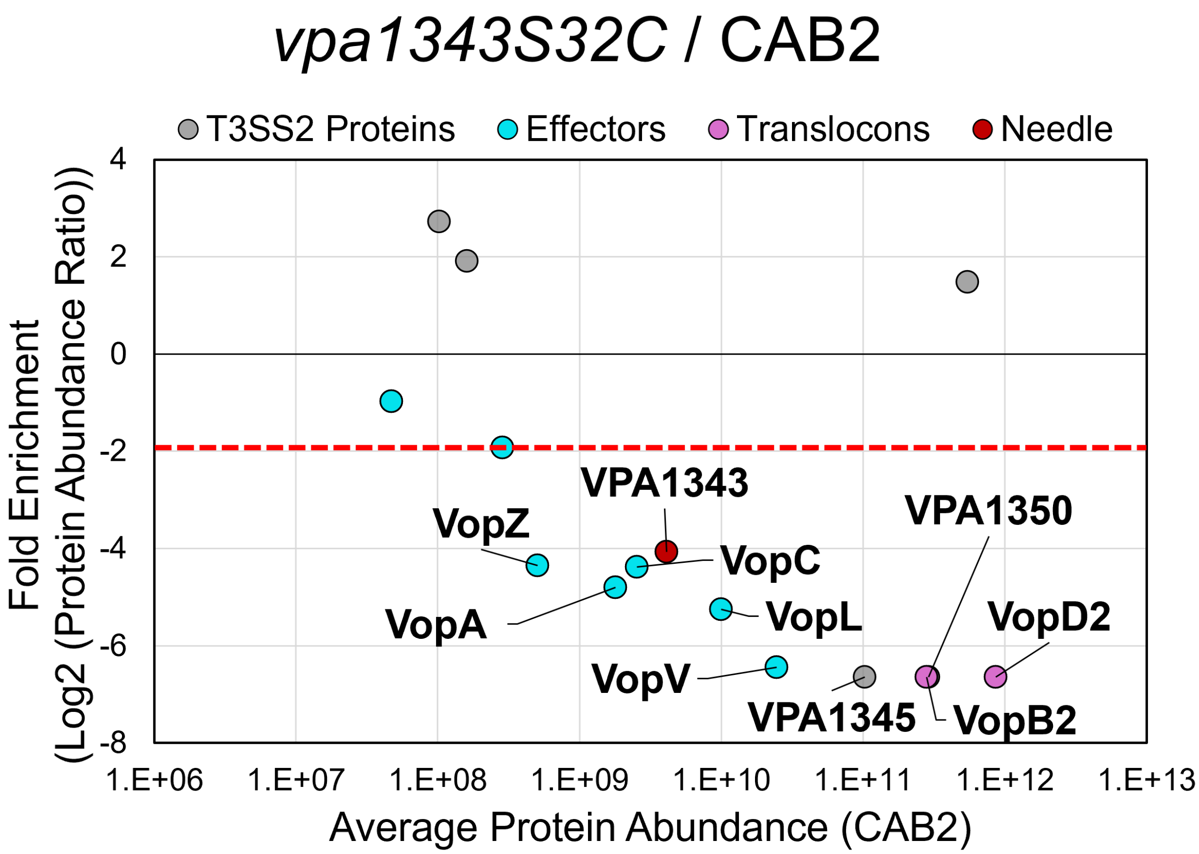

**S5 Fig. Ratio of proteins identified by label-free LC-MS/MS data comparing the CAB2 and *vpa1343S32C* strains.**

Plot of T3SS2 proteins identified by label-free LC-MS/MS comparing *vpa1343S32C and* CAB2 secreted protein fractions. The fold enrichment (*vpa1343*S32C/CAB2) is plotted against the average protein abundance in the CAB2 strain (*n = 3*). The red horizontal dotted line indicates the significance threshold ($Log_2$(*vpa1343*S32C /CAB2) < -2).

**S1 Table. AlphaFold model confidence**

| Chaperone | Amino acid sequence | Partner | Sequence | ipTM | pTM |
|---|---|---|---|---|---|
| VPA1334 | 1-124 | VopC | 1-84 | 0.74 | 0.8 |
| VPA1334 | 1-124 | VopL | 1-78 | 0.73 | 0.79 |
| VPA1352 | 1-137 | VopV | 1-72 | 0.78 | 0.84 |

| | | | | | |
|---|---|---|---|---|---|
| VPA1365 | 1-162 | VopB2 | 1-72 | 0.8 | 0.74 |

**S2 Table. List of different classes of chaperones in well-studied Gram-negative bacteria**

| Bacteria | Class I | | | Class II | | | Class III/V | | |
|---|---|---|---|---|---|---|---|---|---|
| | **Chaperone** | **Substrate (Effectors)** | **Reference** | **Chaperone** | **Substrate (Translocons)** | **Reference** | **Chaperone** | **Substrate (Needle)** | **Reference** |
| *Pseudomonas aeruginosa* | Orf1 | ExoS | (1) | PcrH | PopB, PopD | (2) | PscE, PscG | PscF | (3) |
| EHEC/EPEC | CesT | Tir | (4) | CesD | EspB, EspD | (5) | CesA | EspA | (6) |
| *Yersinia* spp. | SycE<br>SycN<br>SycO<br>SycT | YopE<br>YopN<br>YopO<br>YopT | (7)<br>(8)<br>(9)<br>(10) | LcrH | YopB, YopD | (11) | YscE, YscG | YscF | (12) |
| *Shigella* spp. | Spa15 | IpaA, IpgB1, OspC3 | (13) | IpgC | IpaB, IpaC | (14) | - | - | - |
| *Salmonella* spp. | SicP<br>SigE<br>InvB | SptP<br>SigD<br>SopE, SopE2 | (15)<br>(16)<br>(17) | SscA | SseC | (18) | SsaH, SsaE | SsaG | (19) |
| *Vibrio parahaemolyticus* | VPA1334<br><br>VPA1352 | VopC, VopL<br>VopV | This study | VPA1365 | VopB2 | This study | VPA1340 | VPA1343 | This study |

**S3 Table. List of primers used in the study**

| Primer name | Sequence (5'-3') |
|---|---|
| JJ209 | CAGTCAGCTAGCCGTTTAGCTAATTTACTTCTTCTTG |
| JJ210 | AGAGTTTAGGTTATGAATAAATGAGGTTTAGTCAACTTTTGC |
| JJ211 | AAAAGTTGACTAAACCTCATTTATTCATAACCTAAACTCTTCATC |
| JJ212 | CTGACTCTCGAGTTAAGTGGGGTGGGGATGACCATG |
| JJ230 | CAGTCACTCGAGATGAATAATATCAATCCAGTATATATC |
| JJ231 | CTGACAGAATTCCTAAACCTCATCGAGGATCTCC |
| JJ235 | TAACTCTACATAGATATTAGATTGATTCCTTGATTTATAACTCTTTG |
| JJ236 | TTATAAATCAAGGAATCAATCTAATATCTATGTAGAGTTATTATTTG |
| JJ237 | AGTCTGTCTAGAATTTATTATTTACGTAGCCTACTCG |
| JJ240 | CAGTCAGTCGACATGAGGAGTGTGATACAGAATTATTG |
| JJ296 | TCGAAATGGAGGCTTAATCTACAGA |
| JJ297 | ATCCGCATCTGTAGATTAAGCCTC |
| JJ298 | CAGTCTCAATAACAAGGGGCTGAA |
| JJ299 | CCTTGTTATTGAGACTGCAATAAC |
| JJ342 | GTCACACTCGAGGTACTACAACTTTTACTTTG |
| JJ343 | GACAATCTTTGAGAGAGTAATCAAGGTAAATACATGAATGTT |
| JJ344 | CATTCATGTATTTACCTTGATTACTCTCTCAAAG |
| JJ345 | GACTCAGCTAGCGTGCTAAAAGAAAAAACTTTAATGTTGG |
| JJ360 | CAGTCACTCGAGGTGAGCGAATTTGAAATAGAAG |
| JJ361 | CTGACTGGTACCTCATATCAAATATGGATTTAAC |
| JJ387 | CAGTCAGCTAGCGATTGCATGCTTGCCCAATG |
| JJ388 | TCAAAACGGATGGAAAATAGGCTAGGGTGTTGCTTGGTTA |

| | |
|---|---|
| JJ389 | TAACCAAGCAACACCCTAGCCTATTTTCCATCCGTTTTGAAC |
| JJ390 | CTGACTCTCGAGTTAAAAAGGCTAAAAAATGAAAA |
| JJ435 | CAGTCAGGTACCAATGAGAACACTGCGAGAAATCG |
| JJ436 | CTGACTGAATTCTTATAGTTTCACTTCAGAAGCCAT |
| JJ439 | TCAGTGTCTAGAAATGCTTAAAATTAAACTGCCTC |
| JJ440 | GTCACAGAATTCCTACGATAATTTTGCAGATAGT |
| JJ453 | TCAGTGAGATCTATGAGAACACTGCGAGAAATCGT |
| JJ454 | AGTCTGAAGCTTTTATAGTTTCACTTCAGAAGCCATT |
| JJ455 | TCAGTGAGATCTTTGAAGTTAGGTTCAAAACG |
| JJ456 | TGACTGAAGCTTTCAAGTAGCCACATTGCTCTCA |
| JJ457 | CTGACTGGTACCATTGAAGTTAGGTTCAAAACGG |
| JJ458 | CTGACTGAATTCTCAAGTAGCCACATTGCTCTCA |
| JJ459 | TCAGTGTCTAGAAATGGCGTATAATATTTCGCTGAAC |
| JJ460 | CTGACTGAATTCTGGTTCACGACGACCGTTGAC |
| JJ476 | TCAGTGGGATCCAATGCCAATATTAAATATTAGTAAAT |
| JJ478 | CTGACTGAATTCTTAACCATTTTTCAATAGCACTA |
| JJ489 | CAGTCAGGATCCGGTGAGCGAATTTGAAATAGAAG |
| JJ490 | CTGACTGGTACCTCATATCAAATATGGATTTAAC |
| JJ491 | TGACTGTCTAGAAATGCCAATTAATGTTGTTAATTCTG |
| JJ492 | CTGACTGAATTCTTAACGGTTAGTAACCAAACTTG |
| JJ493 | TGACTGTCTAGAAATGAAAACATCATTAGTGAATA |
| JJ494 | GTCACTGAATTCTTACATATGAATTGAAGTCATT |
| JJ209 | CAGTCAGCTAGCCGTTTAGCTAATTTTACTTCTTCTTG |
| JJ210 | AGAGTTTAGGTTATGAATAAATGAGGTTTAGTCAACTTTTGC |
| JJ211 | AAAAGTTGACTAAACCTCATTTATTCATAACCTAAACTCTTCATC |
| JJ212 | CTGACTCTCGAGTTAAGTGGGGTGGGGATGACCATG |
| JJ230 | CAGTCACTCGAGATGAATAATATCAATCCAGTATATATC |
| JJ231 | CTGACAGAATTCCTAAACCTCATCGAGGATCTCC |
| JJ235 | TAACTCTACATAGATATTAGATTGATTCCTTGATTTATAACTCTTTG |
| JJ236 | TTATAAATCAAGGAATCAATCTAATATCTATGTAGAGTTATTATTTG |
| JJ237 | AGTCTGTCTAGAATTTATTATTTACGTAGCCTACTCG |
| JJ240 | CAGTCAGTCGACATGAGGAGTGTGATACAGAATTATTG |
| JJ296 | TCGAAATGGAGGCTTAATCTACAGA |
| JJ297 | ATCCGCATCTGTAGATTAAGCCTC |
| JJ298 | CAGTCTCAATAACAAGGGGCTGAA |
| JJ299 | CCTTGTTATTGAGACTGCAATAAC |
| JJ342 | GTCACACTCGAGGTACTACAACTTTTACTTTG |
| JJ343 | GACAATCTTTGAGAGAGTAATCAAGGTAAATACATGAATGTT |
| JJ344 | CATTCATGTATTTACCTTGATTACTCTCTCAAAG |
| JJ345 | GACTCAGCTAGCGTGCTAAAAGAAAAAACTTTAATGTTGG |
| JJ360 | CAGTCACTCGAGGTGAGCGAATTTGAAATAGAAG |
| JJ361 | CTGACTGGTACCTCATATCAAATATGGATTTAAC |
| JJ387 | CAGTCAGCTAGCGATTGCATGCTTGCCCAATG |
| JJ388 | TCAAAACGGATGGAAAATAGGCTAGGGTGTTGCTTGGTTA |
| JJ389 | TAACCAAGCAACACCCTAGCCTATTTTCCATCCGTTTTGAAC |

| | |
|---|---|
| JJ390 | CTGACTCTCGAGTTAAAAAGGCTAAAAAATGAAAA |
| JJ435 | CAGTCAGGTACCAATGAGAACACTGCGAGAAATCG |
| JJ436 | CTGACTGAATTCTTATAGTTTCACTTCAGAAGCCAT |
| JJ439 | TCAGTGTCTAGAAATGCTTAAAATTAAACTGCCTC |
| JJ440 | GTCACAGAATTCCTACGATAATTTTGCAGATAGT |
| JJ441 | CAGTCAGCTAGCGATTCAGTATAAATGTGATTTT |
| JJ442 | TCATATGTAAGGAGAAAATTTTAGTTTAAAAGGGAGAGCG |
| JJ443 | CGCTCTCCCTTTTAAACTAAAATTTTCTCCTTACATATGA |
| JJ444 | CTGACACTCGAGACAAAAAGATTCACAAATCGCGA |
| JJ453 | TCAGTGAGATCTATGAGAACACTGCGAGAAATCGT |
| JJ454 | AGTCTGAAGCTTTTATAGTTTCACTTCAGAAGCCATT |
| JJ455 | TCAGTGAGATCTTTGAAGTTAGGTTCAAAACG |
| JJ456 | TGACTGAAGCTTTCAAGTAGCCACATTGCTCTCA |
| JJ457 | CTGACTGGTACCATTGAAGTTAGGTTCAAAACGG |
| JJ458 | CTGACTGAATTCTCAAGTAGCCACATTGCTCTCA |
| JJ459 | TCAGTGTCTAGAAATGGCGTATAATATTTCGCTGAAC |
| JJ460 | CTGACTGAATTCTGGTTCACGACGACCGTTGAC |
| JJ476 | TCAGTGGGATCCAATGCCAATATTAAATATTAGTAAAT |
| JJ478 | CTGACTGAATTCTTAACCATTTTTCAATAGCACTA |
| JJ489 | CAGTCAGGATCCGGTGAGCGAATTTGAAATAGAAG |
| JJ490 | CTGACTGGTACCTCATATCAAATATGGATTTAAC |
| JJ491 | TGACTGTCTAGAAATGCCAATTAATGTTGTTAATTCTG |
| JJ492 | CTGACTGAATTCTTAACGGTTAGTAACCAAACTTG |
| JJ493 | TGACTGTCTAGAAATGAAAACATCATTAGTGAATA |
| JJ494 | GTCACTGAATTCTTACATATGAATTGAAGTCATT |

**S4 Table. List of plasmids and strains used in the study**

| **Plasmids** | | |
|---|---|---|
| **Name** | **Description** | **Source/Reference** |
| pDM4 | Suicide vector, γ ori R6K, sacB, $Cm^R$ | Lab stock |
| pBAD-Fix | Arabinose inducible expression vector suitable for complementation, $Kan^R$ | Lab stock |
| pMW-GFP | Constitutive GFP expressing plasmid, $Spec^R$ | Lab stock |
| pKT25 | Plasmid encoding T25 fragment (amino acids 225-399) of cyaA, $Kan^R$ | Euromedex/ Dr. Zezhang Wen, Louisiana State University Health Sciences Center, New Orleans, Louisiana, USA. |

| | | |
|---|---|---|
| pUT18C | Plasmid encoding T18 fragment (corresponding to first 224 amino acids) of cyaA, $Amp^R$ | Euromedex/ Dr. Zezhang Wen, Louisiana State University Health Sciences Center, New Orleans, Louisiana, USA. |
| pKT25-Zip | pKT25 in which the leucine zipper of GCN4 is genetically fused in frame to the T25 fragment (inserted within the KpnI site of pKT25) | Euromedex/ Dr. Zezhang Wen, Louisiana State University Health Sciences Center, New Orleans, Louisiana, USA. |
| pUT18C-Zip | pUT18C in which the leucine zipper of GCN4 is genetically fused in frame to the T18 fragment (inserted between the KpnI and the EcoRI site of pUT18C) | Euromedex/ Dr. Zezhang Wen, Louisiana State University Health Sciences Center, New Orleans, Louisiana, USA. |
| JJ309 (*vpa1334*±800bp-pDM4) | 800bp flanking region of *vpa1334* cloned in XbaI and SalI site of pDM4, $Cm^R$ | This study |
| ECMT63 (*vpa1352*±800bp-pDM4) | 800bp flanking region of *vpa1352* cloned in NheI and XhoI site of pDM4, $Cm^R$ | This study |
| ECMT42 (*vpa1365*±800bp-pDM4) | 800bp flanking region of *vpa1365* cloned in NheI and XhoI site of pDM4, $Cm^R$ | This study |
| JJ332 (*vpa1361*±800bp-pDM4) | 800bp flanking region of *vpa1361* cloned in NheI and XhoI site of pDM4, $Cm^R$ | This study |
| JJ196 (*vpa1340*±800bp-pDM4) | 800bp flanking region along with coding sequence of *vpa1340* with two stop codons cloned in NheI and XhoI site of pDM4, $Cm^R$ | This study |
| JJ330 (*vpa1334*-pBAD-Fix) | Coding sequence of *vpa1334* cloned in BglII and HindIII site of pBAD-Fix, $Kan^R$ | This study |
| JJ331 (*vpa1352*-pBAD-Fix) | Coding sequence of *vpa1352* cloned in BglII and HindIII site of pBAD-Fix, $Kan^R$ | This study |
| ECMT56 (*vpa1365*-pBAD-Fix) | Coding sequence of *vpa1365* cloned in Xho1 and KpnI site of pBAD-Fix, $Kan^R$ | This study |
| JJ258 (*vpa1340*-pBAD-Fix) | Coding sequence of *vpa1340* cloned in Xho1 and EcoRI site of pBAD-Fix, $Kan^R$ | This study |

| | | |
|---|---|---|
| JJ325 (pKT25-*vpa1334*) | Coding sequence of *vpa1334* cloned in KpnI and EcoRI site of pKT25, Kan$^R$ | This study |
| JJ333 (pKT25-*vpa1352*) | Coding sequence of *vpa1352* cloned in KpnI and EcoRI site of pKT25, Kan$^R$ | This study |
| JJ359 (pKT25-*vpa1365*) | Coding sequence of *vpa1365* cloned in XbaI and KpnI site of pKT25, Kan$^R$ | This study |
| JJ351 (pUT18C-*vopC*) | Coding sequence of *vpa1321* (*vopC*) cloned in BamHI and EcoRI site of pUT18C, Amp$^R$ | This study |
| JJ327 (pUT18C-*vopL*) | Coding sequence of *vpa1370* (*vopL*) cloned in XbaI and EcoRI site of pUT18C, Amp$^R$ | This study |
| JJ334 (pUT18C-*vopV*) | Coding sequence corresponding to first 200 amino acids of *vpa1357* (*vopV*) cloned in XbaI and EcoRI site of pUT18C, Amp$^R$ | This study |
| JJ360 (pUT18C-*vopD2*) | Coding sequence of *vpa1361* (*vopD2*) cloned in XbaI and EcoRI site of pUT18C, Amp$^R$ | This study |
| JJ361 (pUT18C-*vopB2*) | Coding sequence of *vpa1362* (*vopB2*) cloned in XbaI and EcoRI site of pUT18C, Amp$^R$ | This study |
| **Strains** | | |
| ***Vibrio parahaemolyticus*** | | |
| CAB2 | RIMD22106233 Δ*tdh* Δ*trh* Δ*exsA* | Lab stock |
| JJ314 (CAB2Δ*vpa1334*) | CAB2 strain deleted for *vpa1334* | This study |
| JJ304 (CAB2Δ*vpa1352*) | CAB2 strain deleted for *vpa1352* | This study |
| JJ279 (CAB2Δ*vpa1365*) | CAB2 strain deleted for *vpa1365* | This study |
| JJ200 (CAB2*vpa1340**) | CAB2 strain with two stop codons in the coding region of *vpa1340* | This study |
| CAB2*vpa1343S3C* | CAB2Δ*vpa1343*:vpa1343S3C-kpnI | (35) |
| JJ246 (CAB2*vpa1343S3C*: *vpa1340**) | CAB2*vpa1343S3C* strain containing two stop codons in *vpa1340* | This study |
| JJ265 (CAB2*vpa1343S3C*: *vpa1340**+ pBAD-*vpa1340*) | CAB2*vpa1343S3C*: *vpa1340** complemented with pBAD-*vpa1340*, Kan$^R$ | This study |
| CAB2*vpa1343S32C* | CAB2 Δ*vpa1343*::*vpa1343S3C*-kpnI | (35) |

| | | |
|---|---|---|
| CAB2Δ*vopC* | CAB2 strain deleted for *vpa1321* (*vopC*) | Lab stock |
| CAB2Δ*vopL* | CAB2 strain deleted for *vpa1370* (*vopL*) | Lab stock |
| CAB2Δ*vopV* | CAB2 strain deleted for *vpa1357* (*vopV*) | Lab stock |
| JJ363 (CAB2Δ*vopD2*) | CAB2 strain deleted for *vpa1361*(*vopD2)* | This study |
| JJ353 (CAB2Δ*vpa1334* Δ*vpa1352*) | CAB2 strain deleted for *vpa1334* and *vpa1352* | This study |
| JJ342 (CAB2Δ*vpa1334*+pBAD-*vpa1334*) | CAB2Δ*vpa1334* complemented with pBAD-*vpa1334*, $Kan^R$ | This study |
| JJ344 (CAB2Δ*vpa1352*+pBAD-*vpa1352*) | CAB2Δ*vpa1352* complemented with pBAD-*vpa1352*, $Kan^R$ | This study |
| JJ298 (CAB2Δ*vpa1365*+pBAD-*vpa1365*) | CAB2Δ*vpa1365* complemented with pBAD-*vpa1365*, $Kan^R$ | This study |
| JJ217 (CAB2*vpa1340**+pBAD-*vpa1340*) | CAB2*vpa1340** complemented with pBAD-*vpa1340*, $Kan^R$ | This study |
| CAB2-GFP | CAB2 strain transformed with pMW-GFP, $Spec^R$ | Lab stock |
| CAB2Δ*vopC*-GFP | CAB2Δ*vopC* strain transformed with pMW-GFP, $Spec^R$ | Lab stock |
| CAB2Δ*vopL*-GFP | CAB2Δ*vopL* strain transformed with pMW-GFP, $Spec^R$ | Lab stock |
| CAB2Δ*vopV*-GFP | CAB2Δ*vopV* strain transformed with pMW-GFP, $Spec^R$ | Lab stock |
| JJ328 (CAB2Δ*vpa1334*-GFP) | CAB2Δ*vpa1334* strain transformed with pMW-GFP, $Spec^R$ | This study |
| JJ329 (CAB2Δ*vpa1352*-GFP) | CAB2Δ*vpa1352* strain transformed with pMW-GFP, $Spec^R$ | This study |
| JJ233 (CAB2*vpa1340**-GFP) | CAB2*vpa1340**- strain transformed with pMW-GFP, $Spec^R$ | This study |
| JJ347 (CAB2Δ*vpa1334*+pBAD-*vpa1334*-GFP) | CAB2Δ*vpa1334*+pBAD-*vpa1334* strain transformed with pMW-GFP, $Spec^R$ | This study |
| JJ349 (CAB2Δ*vpa1352*+pBAD-*vpa1352*-GFP) | CAB2Δ*vpa1352*+pBAD-*vpa1352* strain transformed with pMW-GFP, $Spec^R$ | This study |

| | | |
|---|---|---|
| JJ233 (CAB2*vpa1340**+pBAD-*vpa1340*-GFP) | CAB2*vpa1340**+pBAD-*vpa1340* strain transformed with pMW-GFP, $Spec^R$ | This study |
| ***Escherichia coli*** | | |
| Mach1 | Host for cloning vectors | Lab stock |
| BTH101 | Reporter strain for bacterial-two hybrid assay | Euromedex/ Dr. Zezhang Wen, Louisiana State University Health Sciences Center, New Orleans, Louisiana, USA. |
| S17 | Host for pDM4 based plasmids suitable for conjugation, λ pir+ | Lab stock |
| DH5α pRK2073 | Strain carrying conjugation helper plasmid pRK2073, $Kan^R$ | Lab stock |
